# Functional genetics with a hypomorphic CENP-C mutant reveals a regulatory system for chromosome congression

**DOI:** 10.1101/2025.08.06.668996

**Authors:** Jiahang Miao, Masatoshi Hara, Kuan-Chung Su, Heather R. Keys, Weixia Kong, Yusuke Takenoshita, Iain M. Cheeseman, Tatsuo Fukagawa

**Author notes:** Correspondence to: Masatoshi Hara or Tatsuo Fukagawa.

## Abstract

Chromosome congression is a key process that acts to align chromosomes at the spindle equator via kinetochore-microtubule interactions, with defects in chromosome alignment leading to chromosomal instability. However, defining the mechanisms that underlie chromosome congression is limited due to the multiple factors that act in parallel to regulate chromosome movement. Here, we conducted a genome-wide Cas9-based functional genetics screen using a hypomorphic CENP-C mutant that affects its kinetochore interactions. Our analysis identified *KIF18A,* whose knockout resulted in synthetic lethality with the CENP-C mutant. Further analysis revealed that the synthetic defect was due to a reduction in CENP-E function in the CENP-C mutant. Our work suggests that KIF18A promotes chromosome alignment in cooperation with CENP-E downstream of CENP-C during early prometaphase. Thus, our analysis enables us to dissect parallel molecular mechanisms for chromosome congression and identify sensitivities and biomarkers that might guide anti-KIF18A chemotherapeutics.

## Introduction

To ensure accurate chromosome segregation during cell division, duplicated chromosomes move to and align at the equator of the mitotic spindle in a process termed chromosome congression.^1-3^ Following nuclear envelope breakdown (NEBD), the two spindle poles separate to form a bipolar mitotic spindle. Concurrently, microtubules from spindle poles capture chromosomes via kinetochores to establish bioriented kinetochore-microtubule attachments, and align chromosomes at the spindle equator. Failure of these processes results in chromosome instability, leading to cell death or aneuploidy, a hallmark of cancer.

The events during chromosome congression are controlled and orchestrated by an intricate regulatory system with multiple factors that function in parallel.^3^ Kinesin and dynein motor proteins regulate microtubule dynamics and bipolar spindle formation.^4,5^ These microtubule motors also provide driving forces for chromosome movement and facilitate kinetochore-microtubule attachments.^6,7^ The kinetochore is a large protein complex that captures and stabilizes spindle microtubules and additionally recruits spindle assembly checkpoint (SAC) proteins to monitor and ensure proper bipolar chromosome attachments.^8,9^ To understand how chromosome congression is achieved, it is crucial to untangle the relationship between the multiple kinetochore-localized regulatory pathways that control chromosome congression. However, many of factors act redundantly to ensure robust chromosome congression, obscuring the roles of these regulators and how they interact functionally.

In vertebrates, the kinetochore has two assembly pathways that act to connect chromosomal DNA with spindle microtubules – mediated by CENP-C and CENP-T.^10^ Both proteins, which are members of inner kinetochore complex termed the constitutive centromere-associated network (CCAN),^10–18^ directly bind to centromeric chromatin and recruit the core outer kinetochore complex, KNL1, Mis12, and Ndc80 complexes (KMN) network (KMN for short), which captures spindle microtubules.^19,20^ However, the individual roles of these two parallel kinetochore assembly pathways remain unclear, including their ability to promote downstream activities such as chromosome congression and segregation.

Although our previous studies demonstrated that the CENP-T pathway generates load-bearing kinetochore-microtubule attachment crucial for chromosome segregation,^21,22^ our recent study indicated that CENP-C supports accurate kinetochore-microtubule attachments.^23^ However, it is still largely unclear what roles CENP-C plays in the parallel pathways that promote chromosome congression and segregation.

To explore the role of the CENP-C pathway in chromosome congression and segregation, we utilized a cell line containing a hypomorphic mutant of *CENP-C*. CENP-C directly associates with the Mis12 complex (Mis12C) of KMN through its Mis12C-binding domain (M12BD).^24–27^ Our previous studies using chicken DT40 and human RPE-1 cells found that, although deleting M12BD from CENP-C (CENP-C^ΔM12BD^) reduces the kinetochore recruitment of KMN and its associated proteins, cells expressing the CENP-C^ΔM12BD^ (CENP-C^ΔM12BD^ cells) grow comparably to control cells despite showing modest mitotic defects.^21,23^ These results indicate that the Mis12C-binding of CENP-C is dispensable for cell growth in vertebrate cultured cells. In addition, since homozygous *CENP-C^ΔM12BD^* mice are viable, the M12BD of CENP-C is dispensable for mouse development.^23^ Given the unique characteristics of *CENP-C^ΔM12BD^*, we hypothesized that CENP-C^ΔM12BD^ would enable us to decipher the roles of the two kinetochore assembly pathways and their relationship.

In this study, we conducted a genome-wide CRISPR/Cas9-based functional genetics screen with human CENP-C^ΔM12BD^ cells to reveal genes that are essential for cell growth in CENP-C^ΔM12BD^ cells but not in control cells. Our screening revealed a synthetic lethality between *CENP-C^ΔM12BD^*and knockouts for *KIF18A*, encoding a plus-end-directed motor. KIF18A controls microtubule dynamics and suppresses the oscillatory movement of chromosomes with end-on attachments, leading to chromosome alignment at the equator in later prometaphase and metaphase.^28–31^ Further analysis of KIF18A in near-diploid human RPE-1 cells expressing CENP-C^ΔM12BD^ revealed that the synthetic defect could be attributed to the reduction in the kinetochore localization of CENP-E, another plus-end-directed motor.^32–34^ Given that CENP-E is crucial for the congression of peripheral chromosomes, which reside near one of the spindle poles without the end-on attachment after NEBD, our findings unmask a function for KIF18A, which cooperates with CENP-E downstream of CENP-C for peripheral chromosome congression during early prometaphase. Thus, this strategy enables us to dissect the molecular mechanisms that underlie the elaborate regulatory system for chromosome congression and consequent faithful chromosome segregation during mitosis.

## Results

### A genome-wide CRISPR/Cas9 screen identifies genes whose knockout exhibits synthetic lethality with *CENP-C^ΔM12BD^*

To understand the components that constitute the regulatory system for chromosome congression and segregation, we conducted a genome-wide CRISPR/Cas9-based synthetic lethal screen with a *CENP-C* mutant lacking its Mis12-binding domain (M12BD) (Figure 1A).^21,23,24,27,35,36^ Our previous studies found that the deletion of M12BD from CENP-C (CENP-C^ΔM12BD^) reduced the levels of the kinetochore-localized outer kinetochore KMN network and their associated proteins in chicken DT40 and human RPE-1 cells. However, cells expressing CENP-C^ΔM12BD^ are viable and grow comparable to control cells.^21,23^

**Figure 1.**
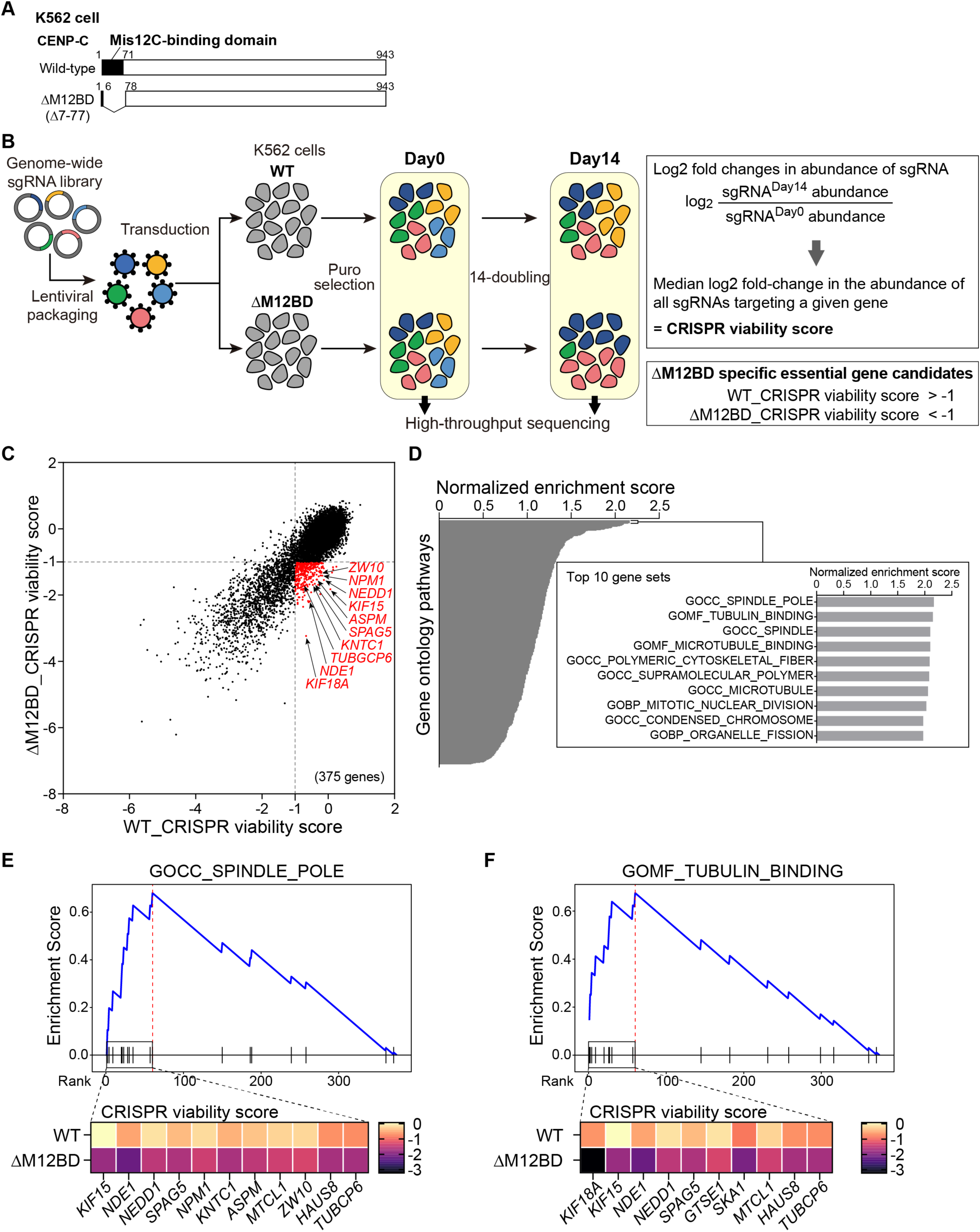
CRISPR screening to identify genes whose knockout exhibits synthetic lethality with *CENP-C^ΔM12BD^* mutation. (A) Schematic representation of human CENP-C protein. The Mis12-binding domain (M12BD) of human CENP-C is highlighted in wild-type (WT) CENP-C. To establish cells expressing CENP-C lacking M12BD (CENP-C^ΔM12BD^ cells), using CRISPR/Cas9 genome editing, exon2-4 encoding amino acids 7-77 were deleted from the *CENP-C* gene locus (see also Figure S1). (B) Schematic representation of the pooled CRISPR screening in K562 WT or CENP-C^ΔM12BD^ cells. Using CRISPR viability scores, genes which are essential for cell growth in K562 CENP-C^ΔM12BD^ cells but not in K562 WT cells were selected as candidates whose knockout showed synthetic lethality with *CENP-C*^ΔM12BD^. (C) Scatter plots showing the CRISPR viability score in K562 WT versus CENP-C^ΔM12BD^ cell pools. Red dots are essential genes in K562 CENP-C^ΔM12BD^ cells but not in K562 WT cells. All CRISPR viability scores in K562 WT and CENP-C^ΔM12BD^ cells are shown in Table S1. (D) Gene set enrichment analysis (GSEA) for 375 candidate genes. The candidate genes were ranked by the magnitude of differences in CRISPR viability score between K562 WT and CENP-C**^Δ^**^M12BD^ cells in descending order and applied to GSEA. The top 10 enriched gene sets were highlighted. (E) GSEA result of GOCC_SPINDLE_POLE gene set. The heat map indicates the CRISPR viability score of the leading edge genes of the preranking based on the magnitude of differences in CRISPR viability score between K562 WT and CENP-C**^Δ^**^M12BD^ cells, in descending order. (F) GSEA result of GOMF_TUBULIN_BINDING gene set. The heat map indicates the CRISPR viability score of the leading edge genes as in (E).

To utilize a genome-wide CRISPR screening system using leukemia-derived K562 cells, we first generated K562 cells expressing CENP-C^ΔM12BD^ by deleting exons 2-4 from *CENP-C* gene, which resulted in the deletion of amino acids (aa) 7-77 encoding most of the M12BD sequence (Figures 1A, S1A-S1D). Similar to the RPE-1 cells used in our previous study,^23^ K562 cells expressing CENP-C^ΔM12BD^ (K562 CENP-C^ΔM12BD^ cells) were viable but displayed reduced kinetochore-localized KMN subcomplexes and a slight increase in mitotic defects (Figures S2A-S2H). Importantly, the cell growth rate of K562 CENP-C^ΔM12BD^ cells was comparable to that of control K562 wild-type cells (K562 WT cells) (Figure S2A).

Using K562 WT and CENP-C^ΔM12BD^ cells, we performed pooled genome-wide CRISPR/Cas9-based functional genetics screening (Figure 1B).^37^ K562 WT or CENP-C^ΔM12BD^ cells were transduced with a lentivirus single guide RNA (sgRNA) knockout library ^38^ targeting 18,663 genes and cultured for 14 population doublings (Figure 1B). Deep sequencing of the sgRNA representation allowed us to calculate a CRISPR viability score reflecting cell viability and proliferation after sgRNA targeting for each gene (Figures 1B, S3A and S3B, and Table S1). A negative viability score indicates that the knockout of the corresponding gene results in reduced cell viability, and genes with scores less than –1 are defined as essential genes under a given growth condition (Figures S3A and S3B).

Targeting most genes resulted in similar overall fitness effects in both K562 WT and CENP-C^ΔM12BD^ cells (Figures 1C, S3A and S3B), with *R^2^*of 0.79 for K562 WT versus K562 CENP-C^ΔM12BD^ cells (Figure 1C). However, a subset of gene knockouts displayed reduced viability scores in K562 CENP-C^ΔM12BD^ cells compared to those in K562 WT cells (Figure 1C). This includes a total of 375 genes (Figure 1C; red dots). Gene set enrichment analysis (GSEA) ^39,40^ found that spindle pole, tubulin-binding, spindle, microtubule-binding, polymeric cytoskeletal fiber, supramolecular polymer, and microtubule-related gene sets were enriched amongst the depleted genes (Figure 1D). Several genes were commonly found in the core enrichment gene list of the gene sets (Figures 1E, 1F, S3C-S3F). Based on their relative viability scores in K562 WT and CENP-C^ΔM12BD^ cells, we selected 10 genes for further validation: *KIF18A*, *KIF15*, *NDE1*, *NEDD1*, *SPAG5*, *NPM1*, *KNTC1*, *ASPM*, *ZW10,* and *TUBGCP6*.

To verify the CRISPR screening results, we conditionally knocked out each of the selected genes in K562 WT or CENP-C^ΔM12BD^ cells. For these experiments, we integrated a doxycycline (Dox)-inducible *Cas9* and three different sgRNAs for each target gene in the cells (Figures S4A-S4C). All genes we tested showed a reduction of their transcript levels after Dox addition (Figures S4D-S4M). We note that after DOX addition, the protein levels of the tested genes were also reduced, except for those of KNTC1 and ASPM under the condition. As shown in Figure 2A, knockout of either one of six genes (*KIF18A*, *KIF15*, *NDE1*, *NEDD1*, *SPAG5,* or *NPM1*) significantly reduced viability in K562 CENP-C^ΔM12BD^ cells relative to K562 WT cells at 4 days after Dox addition. *NEDD1* knockout substantially reduced the viability in K562 CENP-C^ΔMBD^ cells but also lowered viability in K562 WT cells (Figure 2A). Although the knockout of two genes (*NDE1* and *NPM1*) reduced the viability of K562 CENP-C^ΔM12BD^ cells compared to K562 WT cells, the changes were modest (Figure 2A). Knockout of the remaining genes (*KNTC1*, *ASPM*, or *ZW10*) did not show changes in viability in K562 WT or CENP-C^ΔM12BD^ cells in this assay (Figure 2A). Further microscopic observation found that knockout of *KIF18A*, *KIF15*, *NDE1*, *NEDD1*, or *SPAG5* increased the mitotic population in K562 CENP-C^ΔM12BD^ cells relative to K562 WT cells (Figures 2B and 2C). In addition, knockout of *KIF18A*, *KIF15*, *NEDD1*, *SPAG5*, *KNTC1*, *ASPM*, or *TUBGCP1* increased cells with micronuclei in K562 CENP-C^ΔM12BD^ cells more than in K562 WT cells (Figure 2C). These results confirm that our large-scale functional genetics screen successfully isolated genes that exhibit synthetic mitotic defects in cells containing the *CENP-C^ΔM12BD^* mutant.

**Figure 2.**
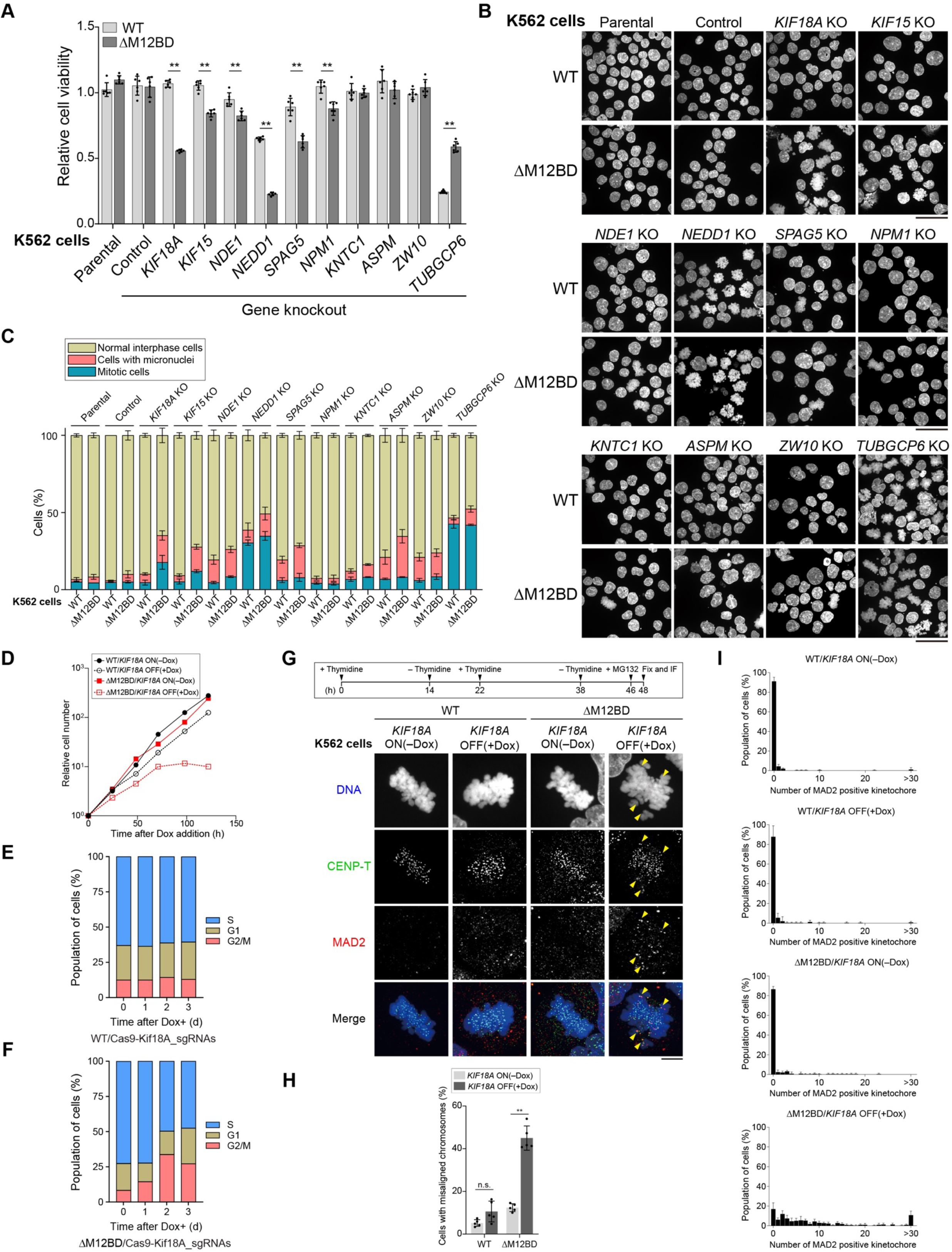
Characterization of phenotype in K562 WT or CENP-C^ΔM12BD^ cells after knockout of genes identified by the CRISPR screening. (A) Cell viability of K562 WT or CENP-C^ΔM12BD^ cells after knockout of indicated genes at 4 days after doxycycline (Dox) addition (Mean and SD, two-tailed Student’s t-test, CENP-C^WT^ cells: n = 6, CENP-C^ΔM12BD^ cells: n = 6; \*\**p* < 0.01). (B) Representative images of DAPI-stained K562 WT or CENP-C^ΔM12BD^ cells after knockout of indicated genes at 4 days. Scale bar, 50 μm. (C) Population of normal interphase cells, mitotic cells, and cells with micronuclei in K562 WT or CENP-C^ΔM12BD^ cells after knockout of indicated genes at 4 days. Error bars indicated SEM. n = 3 independent experiments; 200 cells from each cell line were quantified in each experiment. (D) The growth curve of K562 WT or CENP-C^ΔM12BD^ cells with or without *KIF18A* knockout. K562 WT or CENP-C^ΔM12BD^ cells were treated with or without Dox (*KIF18A* OFF or ON). The cell numbers were normalized to those at time 0 of each line. Dox was added at the time 0. (E) Cell-cycle distribution of conditional knockout of *KIF18A* in K562 WT cells at each day after Dox addition, based on FACS analysis. (F) Cell-cycle distribution of conditional knockout of *KIF18A* in K562 CENP-C^ΔM12BD^ cells at each day after Dox addition, based on FACS analysis. (G and H) Quantification of cells with misaligned chromosomes in K562 WT or CENP-C^ΔM12BD^ cells with or without *KIF18A* knockout. The cells were treated with thymidine for 14 h. Then, thymidine was washed out and incubated for 8 h. Thymidine was added again and cells were incubated for 16 h. After washing thymidine out and incubating for 8 h, MG132 was added and incubated for 2 h (–Dox, *KIF18A* ON). For *KIF18A* knockout, the cells were treated and cultured in the presence of Dox (+Dox, *KIF18A* OFF). The treated cells were fixed and stained with antibodies against MAD2 (red) to detect misaligned chromosomes and CENP-T (green) as a kinetochore marker. DNA was stained with DAPI (blue). Arrowheads show typical MAD2 positive unaligned chromosomes. Scale bar, 10 μm. The cells with MAD2 positive chromosomes were quantified (F). (Mean and SEM, two-tailed Student’s t-test, n = 5 independent experiments; n.s., non-significant; \*\**p* < 0.01). (I) Numbers of MAD2 positive kinetochores in each cell in each condition (WT *KIF18A* ON; WT *KIF18A* OFF; CENP-C^ΔM12BD^ *KIF18A* ON; CENP-C^ΔM12BD^ *KIF18A* OFF) (Mean and SEM, n = 5 independent experiments).

### *KIF18A* knockout with *CENP-C^ΔM12BD^* results in chromosome alignment errors

Among the genes we tested, *KIF18A* knockout showed the most evident synthetic mitotic defects with *CENP-C^ΔM12BD^* in K562 cells. In addition, *KIF18A* conditional knockout cells showed a significant reduction in cell viability even at 2 days after Dox addition (Figure S4N). Therefore, we examined the synergistic effects of *KIF18A* knockout in depth in K562 CENP-C^ΔM12BD^ cells.

We first examined the cell proliferation rate of K562 CENP-C^ΔM12BD^ cells after *KIF18A* knockout (Figure 2D). Consistent with the cell viability assay, *KIF18A* knockout caused a growth arrest and subsequent cell death in K562 CENP-C^ΔM12BD^ cells (*ΔM12BD/KIF18A* OFF, Figure 2D) but not in K562 WT cells (*WT/KIF18A* OFF, Figure 2D). Based on flow cytometry analysis, the cell-cycle distribution did not display significant changes before and after Dox addition in K562 WT cells but showed an increased G2/M population in K562 CENP-C^ΔM12BD^ cells after Dox addition (Figures 2E and 2F). This suggests that *KIF18A* knockout caused a mitotic arrest in K562 CENP-C^ΔM12BD^ cells. To further test the mitotic arrest, we conducted immunofluorescence with an antibody against MAD2, a SAC protein that localizes to misaligned chromosomes. We found that the *KIF18A* knockout cells showed significantly increased misaligned chromosomes in K562 CENP-C^ΔM12BD^ but not in WT cells (Figures 2G-2I). This suggests that the *KIF18A* knockout results in the chromosome alignment defects and subsequent cell death in K562 CENP-C^ΔM12BD^ cells.

### KIF18A is required for cell viability in RPE-1 CENP-C^ΔM12BD^ cells

Based on the mitotic defect and mitotic arrest observed in *KIF18A* knockout + CENP-C^ΔM12BD^ K562 cells, we next sought to test whether a similar synthetic lethality occurs in other cell types. To do this, we tested the near-diploid human RPE-1 cell line, which is insensitive to *KIF18A* knockdown.^41^ Our previous work found that RPE-1 CENP-C^ΔM12BD^ cells displayed similar cell proliferation to control RPE-1 cells (Figures 3A, S5A-G).^23^ To test cell viability after *KIF18A* knockdown, we treated RPE-1 CENP-C^WT^ or CENP-C^ΔM12BD^ cells with siRNAs targeting *KIF18A* (siKIF18A). KIF18A protein was undetectable after 24 h in the cells treated with siKIF18A, but not with a control non-targeting siRNA (siNT) (Figure 3B). Although siKIF18A treatment did not cause growth defects in RPE-1 CENP-C^WT^ cells, viability was significantly reduced in RPE-1 CENP-C^ΔM12BD^ cells treated with siKIF18A (Figure 3C). This indicates that the deletion of the CENP-C Mis12 complex binding domain leads to *KIF18A* knockdown sensitivity (KIF18A-KD sensitivity) in RPE-1 cells.

**Figure 3.**
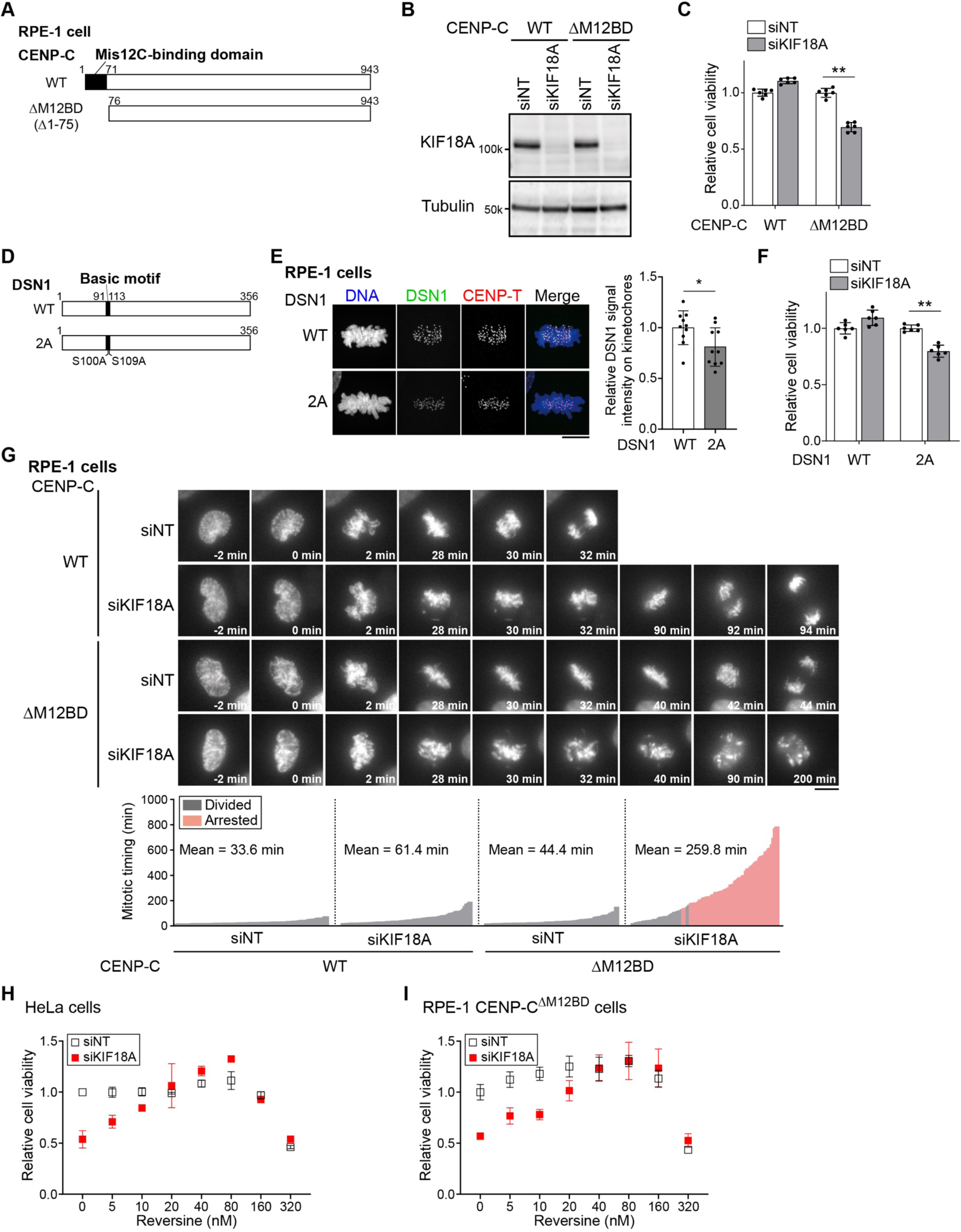
Abnormal mitotic progression in RPE-1 CENP-C^ΔM12BD^ cells after knockdown of KIF18A. (A) Schematic representation of human wild-type CENP-C (CENP-C^WT^) and CENP-C^ΔM12BD^ mutant, which are expressed in RPE-1 cells. To generate RPE-1 CENP-C^ΔM12BD^ cells, *CENP-C* cDNA encoding full length (WT: aa 1-943) or aa 76-943 (ΔM12BD: Δaa 1-75) was introduced into the endogenous *CENP-C* locus, respectively (See also Figure S5). (B) Immunoblot analysis to detect KIF18A in RPE-1 CENP-C^WT^ or CENP-C^ΔM12BD^ cells after treatment with control Non-Targeting siRNA (siNT) or KIF18A siRNA (siKIF18A). α-tubulin was probed as a loading control. (C) Cell viability assay of RPE-1 CENP-C^WT^ or CENP-C^ΔM12BD^ cells after treatment with siNT or siKIF18A. (Mean and SD, two-tailed Student’s t-test, CENP-C^WT^ cells with siNT: n = 6, CENP-C^WT^ cells with siKIF18A: n = 6, CENP-C^ΔM12BD^ cells with siNT: n = 6, CENP-C^ΔM12BD^ cells with siKIF18A: n = 6; \*\**p* < 0.01). (D) Schematic representation of human wild-type DSN1 (DSN1^WT^) and DSN1^2A^ (S100A/S109A) mutant, which are expressed in RPE-1 cells. DSN1 has two Aurora B phosphorylation sites at S100 and S109. These two Ser residues are replaced with Ala in DSN1^2A^ mutant. To generate RPE-1 DSN1^WT^ or DSN1^2A^ cells, *mScarlet*-fused *DSN1* cDNA encoding DSN1^WT^ or DSN1^2A^ was introduced into the endogenous *DSN1* locus, respectively (See also Figure S5). (E) mScarlet-DSN1 localization in RPE-1 DSN1^WT^ or DSN1^2A^ cells. mScarlet-DSN1 is shown in green. CENP-T was stained as a kinetochore marker (CENP-T, red), DNA was stained with DAPI (blue). Scale bar, 10 μm. Signal intensities of mScarlet-DSN1 at mitotic kinetochores were quantified (Mean and SD, two-tailed Student’s t-test, DSN1^WT^ cells: n = 10 cells, DSN1^2A^ cells: n = 10 cells; \**p* < 0.1). (F) Cell viability of RPE-1 DSN1^WT^ or DSN1^2A^ cells after treatment with siNT or siKIF18A. (Mean and SD, two-tailed Student’s t-test, DSN1^WT^ cells with siNT: n = 6, DSN1^WT^ cells with siKIF18A: n = 6, DSN1^2A^ cells with siNT: n = 6, DSN1^2A^ cells with siKIF18A: n = 6; \*\**p* < 0.01,). (G) Representative time-lapse images of mitotic progression in RPE-1 CENP-C^WT^ or CENP-C^ΔM12BD^ cells after treatment with siNT or siKIF18A. DNA was visualized with GFP-H2A. Images were projected using maximum intensity projection. Time is relative to nuclear envelope breakdown (NEBD). Scale bar, 10 μm. The time-lapse images were analyzed to measure the mitotic timing from NEBD to anaphase onset (CENP-C^WT^ cells with siNT: n = 99 cells, CENP-C^WT^ cells with siKIF18A: n = 84 cells, CENP-C^ΔM12BD^ cells with siNT: n = 86 cells, CENP-C^ΔM12BD^ cells with siKIF18A: n = 95 cells). The red bars show the cells that were arrested in mitosis and did not exit mitosis by the end of the imaging. (H) Cell viability of HeLa cells in the presence of various concentrations of Reversine after treatment of siNT or siKIF18A (Mean and SD, two-tailed Student’s t-test, each sample size: n = 3). (I) Cell viability of RPE-1 CENP-C^ΔM12BD^ cells in the presence of various concentrations of Reversine after treatment with siNT or siKIF18A (Mean and SD, two-tailed Student’s t-test, n = 6).

We next sought to determine whether the KIF18A-KD sensitivity in RPE-1 CENP-C^ΔM12BD^ cells depends on altered Mis12C-CENP-C interactions or an alternative role of the M12BD in CENP-C. CENP-C-Mis12C binding is facilitated by Aurora B phosphorylation of DSN1, a Mis12C subunit, at Ser100 and Ser109.^25,26^ In DSN1 phospho-deficient mutants (DSN1 S100A/S109A, DSN1^2A^), the binding of Mis12C to CENP-C is reduced (Figure S5H).^27,42,43^ We introduced DSN1^WT^ or DSN1^2A^ into the endogenous *DSN1* locus in RPE-1 cells (RPE-1 DSN1^WT^ or DSN1^2A^ cells, Figures 3D, S5I and S5J). As reported previously,^25^ DSN1^2A^ reduced the centromeric localization of Mis12C (DSN1) as well as Ndc80C (HEC1), KNL1C (KNL1), BubR1, and ZW10 in RPE-1 cells (Figures 3E, S5K-S5N). siKIF18A treatment significantly reduced the cell viability of RPE-1 DSN1^2A^, but not DSN1^WT^ cells (Figure 3F). However, the reduction of viability in RPE-1 DSN1^2A^ cells after *KIF18A* knockdown was milder than that in RPE-1 CENP-C^ΔM12BD^ cells, possibly due to less reduction of Mis12C localization in RPE-1 DSN1^2A^ cells (Figures 3E and S5D). Nevertheless, our results suggest that perturbating Mis12C-CENP-C interactions leads to increased KIF18A-KD sensitivity.

As we observed chromosome alignment errors in K562 CENP-C^ΔM12BD^ cells after *KIF18A* knockout (Figures 2G-2I), we next examined mitotic progression by time-lapse imaging in RPE-1 CENP-C^WT^ or CENP-C^ΔM12BD^ cells treated with control siNT or siKIF18A. RPE-1 CENP-C^ΔM12BD^ cells showed a slight delay in mitotic progression from nuclear envelope breakdown (NEBD) to anaphase onset in the presence of siNT (siNT RPE-1 CENP-C^WT^ cells: 33 min, siNT RPE-1 CENP-C^ΔM12BD^ cells: 44 min, Figure 3G, Movies S1 and S3), consistent with our previous study.^23^ Although the *KIF18A* knockdown caused a slight mitotic delay in RPE-1 CENP-C^WT^ cells (61 min), the observed cells exited from mitosis (Figure 3G and Movie S2). In contrast, the *KIF18A* knockdown in CENP-C^ΔM12BD^ cells resulted in severe chromosome alignment defects and mitotic arrest in 63% of cells during the observation (Figure 3G and Movie S4). These results indicate that the deletion of the CENP-C M12BD combined with *KIF18A* knockdown causes a synthetic mitotic defect and prolonged mitotic arrest in RPE-1 cells. Thus, RPE-1 CENP-C^ΔM12BD^ cells require KIF18A for their mitotic progression and cell viability.

A previous study found that one of the determinants for sensitivity to KIF18A inhibition in cancer cells was a prolonged mitosis but that SAC perturbation suppressed KIF18A sensitivity in those cells.^41^ In line with this previous study, *KIF18A* knockdown reduced cell viability in HeLa cells, a cancer cell line sensitive to KIF18A inhibition, and this reduction was suppressed by SAC inhibition using low doses of the Mps1 inhibitor Reversine (Figure 3H). Reversine treatment also suppressed the reduced viability observed after siKIF18A treatment in RPE-1 CENP-C^ΔM12BD^ cells (Figure 3I), suggesting that a prolonged mitotic arrest is at least partially responsible for the KIF18A-KD sensitivity in RPE-1 CENP-C^ΔM12BD^ cells.

### KIF18A microtubule plus-end localization is required for cell viability in CENP-C^ΔM12BD^ cells

KIF18A is a plus-end directed kinesin that accumulates at the plus-end tips of microtubules and suppresses kinetochore-microtubule dynamics.^28,29^ To understand how KIF18A is required for RPE-1 CENP-C^ΔM12BD^ cell viability, we dissected the functional domains in KIF18A. We integrated siRNA-resistant WT or mutants *KIF18A* cDNAs at the *AAVS1* locus to stably express them in RPE-1 CENP-C^WT^ or CENP-C^ΔM12BD^ cells and examined their viability after siNT or siKIF18A treatment (Figures 4A-E and S6). In control RPE-1 CENP-C^WT^ cells, RNAi-based replacements with these *KIF18A* transgenes did not reduce cell viability (Figure 4D). Similarly, in RPE-1 CENP-C^ΔM12BD^ cells, replacement with a KIF18A^WT^ construct did not alter cell viability. In contrast, replacement with a KIF18A mutant lacking the motor domain, which is required for KIF18A localization to the plus-end tips of kinetochore microtubules,^28^ resulted in reduced viability similar to KIF18A-depleted cell (Figures 4E and 4F).

**Figure 4.**
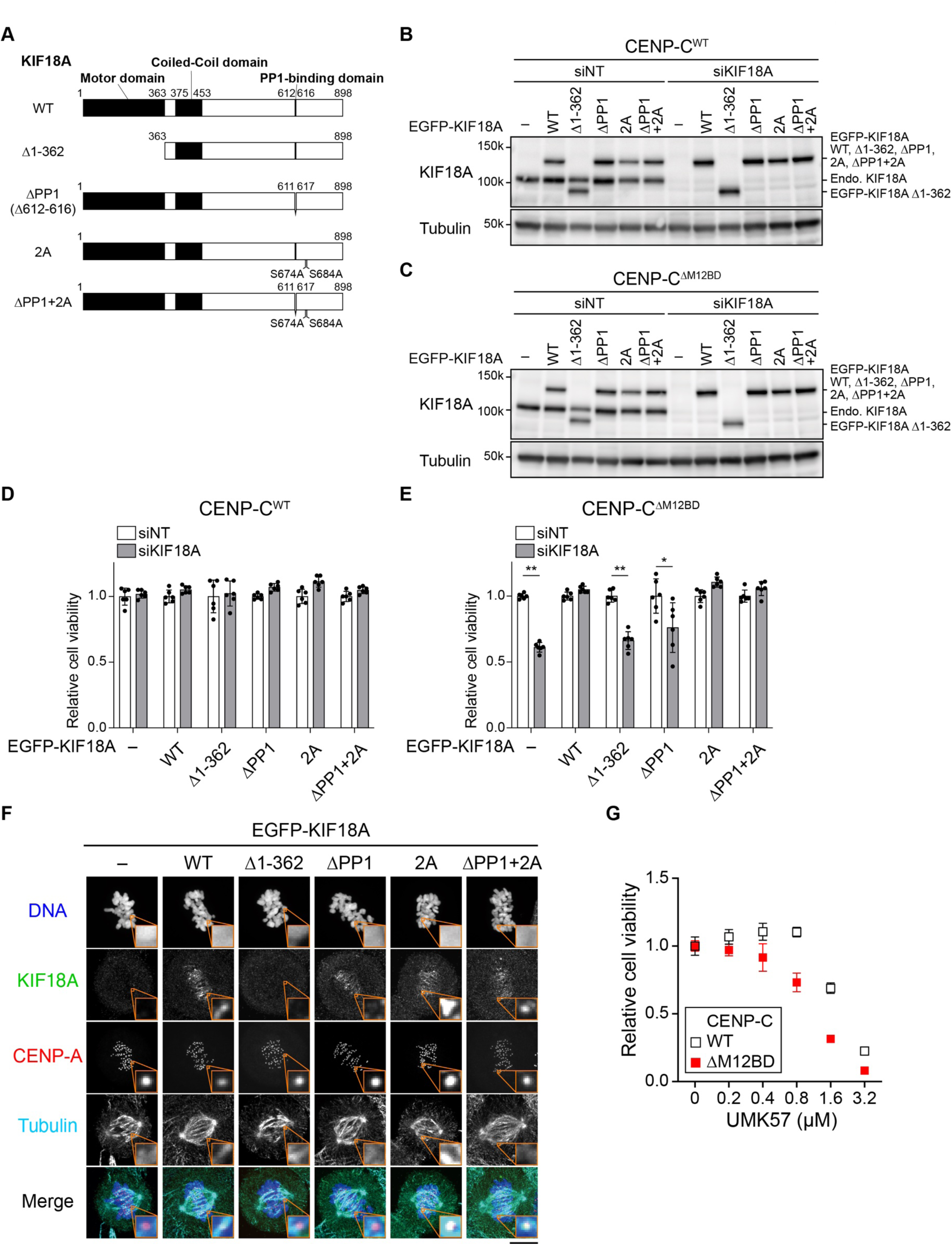
Microtubule plus-end localization of KIF18A is required for cell viability in RPE-1 CENP-C^ΔM12BD^ cells. (A) Schematic representation of wild-type KIF18A and its functional domains, including the motor domain, coiled-coil domain, and PP1 binding domain. RPE-1 cells expressing KIF18A lacking the motor domain (KIF18A^Δ1-362^) or PP1 binding domain (KIF18A^ΔPP1^) were generated. KIF18A is phosphorylated by CDK1 at S674 and S684. These phosphorylation sites are dephosphorylated by PP1 bound to KIF18A. RPE-1 cells expressing a KIF18A mutant in which S674 and S684 were replaced with Ala (KIF18A^2A^) or KIF18A^2A^ lacking PP1 binding domain (KIF18A^ΔPP1+2A^) were also generated (also see Figure S6). (B and C) Immunoblot analysis to detect WT or mutant KIF18A protein in RPE-1 CENP-C^WT^ (B) or CENP-C^ΔM12BD^ (C) cells with siNT or siKIF18A treatment. α-tubulin was probed as a loading control. (D and E) Cell viability assay of RPE-1 CENP-C^WT^ (D) or CENP-C^ΔM12BD^ (E) cells expressing WT or various KIF18A mutants with siNT or siKIF18A treatment (Mean and SD, two-tailed Student’s t-test, each sample size: n = 6; \**p* < 0.1; \*\**p* < 0.01). (F) Localization of KIF18A^WT^ and its mutants (green) at plus-end of microtubules. The cells were treated with siKIF18A for 2 days before fixation. KIF18A was stained with an antibody against human KIF18A. mScarlet-CENP-A was used as a kinetochore marker (red) and microtubules were stained with an anti-α-tubulin antibody (cyan). DNA was stained with DAPI (blue). Scale bar, 10 μm. (G) Cell viability assay of RPE-1 CENP-C^WT^ or CENP-C^ΔM12BD^ cells with UMK57 at various concentrations. (Mean and SD, two-tailed Student’s t-test, each sample size: n = 6).

KIF18A also has a domain that interacts with Protein phosphatase 1 (PP1) (PP1-binding domain, Figure 4A).^44^ KIF18A-PP1 interactions are required for KIF18A localization to the microtubule plus-end tips by dephosphorylating KIF18A at two CDK1 phosphorylation sites, Ser674 and Ser684.^44^ Deletion of the PP1-binding domain (ΔPP1) prevents KIF18A from localizing to microtubule plus ends but the addition of alanine substitutions at Ser674/Ser684 (ΔPP1+2A) restores the localization of KIF18A lacking the PP1-binding domain (KIF18A^ΔPP1^ and KIF18A^ΔPP1+2A^, Figure 4F).^44^ Consistent with this localization, KIF18A^ΔPP1+2A^, but not KIF18A^ΔPP1^, rescued the reduction of cell viability caused by *KIF18A* knockdown in RPE-1 CENP-C^ΔM12BD^ cells. These results demonstrate that the plus-end tip localization of KIF18A is required for cell viability in RPE-1 CENP-C^ΔM12BD^ cells.

KIF18A suppresses kinetochore-microtubule dynamics, stabilizing kinetochore-microtubule attachments.^28,29,45^ To test whether the regulation of kinetochore-microtubule dynamics is critical for the viability of RPE-1 CENP-C^ΔM12BD^ cells, we treated the cells with the KIF2C/MCAK agonist UMK57, which destabilizes kinetochore microtubules.^46^ Sublethal doses of UMK57 reduced the cell viability of RPE-1 CENP-C^ΔM12BD^ cells (0.4-0.8 μM; Figure 4G), suggesting that RPE-1 CENP-C^ΔM12BD^ cells require stable kinetochore microtubules. Supporting this result, our functional genetic screen reveals that knockouts of the MCAK suppressor GTSE1^47^ also display synthetic lethality with CENP-C ΔM12BD mutation (Figures 1F and S3D-S3F and Table S1).

Next, we investigated the source of the KIF18A-KD sensitivity in RPE-1 CENP-C^ΔM12BD^ cells. We tested whether the reduction of Ndc80C sensitized RPE-1 cells to *KIF18A* knockdown. For this, we used cells expressing CENP-T mutants that lack either one of the two Ndc80C-binding domains (Figures S7A and S7B).^23^ However, in contrast to conditions with reduced Mis12 localization, the viability of the cells expressing CENP-T mutants remained unchanged after siKIF18A treatment (Figure S7C). We also tested the effect of Aurora B reduction on the KIF18A-KD sensitivity, as Aurora B levels were significantly reduced in RPE-1 CENP-C^ΔM12BD^ cells.^23^ siKIF18A treatment did not cause obvious changes in the cell viability of RPE-1 cells treated with sublethal doses of the Aurora B inhibitor AZD1152 (Figure S7D). These results suggest that neither Ndc80C nor Aurora B reduction in RPE-1 CENP-C^ΔM12BD^ cells is directly responsible for mitotic defects with *KIF18A* knockdown, indicating that there must be an alternate cause of the KIF18A-KD sensitivity in RPE-1 CENP-C^ΔM12BD^ cells.

### Reduced CENP-E kinetochore localization explains the *KIF18A*-KD sensitivity of CENP-C^ΔM12BD^ cells

To seek the cause of the KIF18A-KD sensitivity in RPE-1 CENP-C^ΔM12BD^ cells, we closely inspected the time-lapse images for mitotic progression (Figure 3G and Movies S1-S4). In the time-lapse images, the siKIF18A-treatment caused prolonged mitotic arrest, which consequently collapsed the mitotic apparatus due to kinetochore-microtubule detachment or premature separation of sister chromatids, known as cohesion fatigue.^48^ Given the SAC inactivation suppressed the reduced viability of the siKIF18A-treated RPE-1 CENP-C^ΔM12BD^ cells (Figure 3I), siKIF18A-treatment could cause defects in chromosome dynamics, leading to prolong mitotic arrest. To test the idea, we looked at chromosome behaviors during early time points after NEBD.

More than 90% of control siNT-treated RPE-1 CENP-C^WT^ cells aligned their chromosomes at the spindle equator within 25 min after NEBD (Figure 5A). In contrast, approximately 50% of siKIF18A-treated RPE-1 CENP-C^WT^ cells still showed unaligned chromosomes at this time point, and it took more than 70 min after NEBD for 90% of cells to fully align their chromosomes (Figure 5A). RPE-1 CENP-C^ΔM12BD^ cells treated with control siNT also showed delayed chromosome congression, compared with CENP-C^WT^ cells (Figures 5A). In siKIF18A-treated RPE-1 CENP-C^ΔM12BD^ cells, the unaligned chromosomes that emerged after NEBD were sustained during observation (Figure 5A).

**Figure 5.**
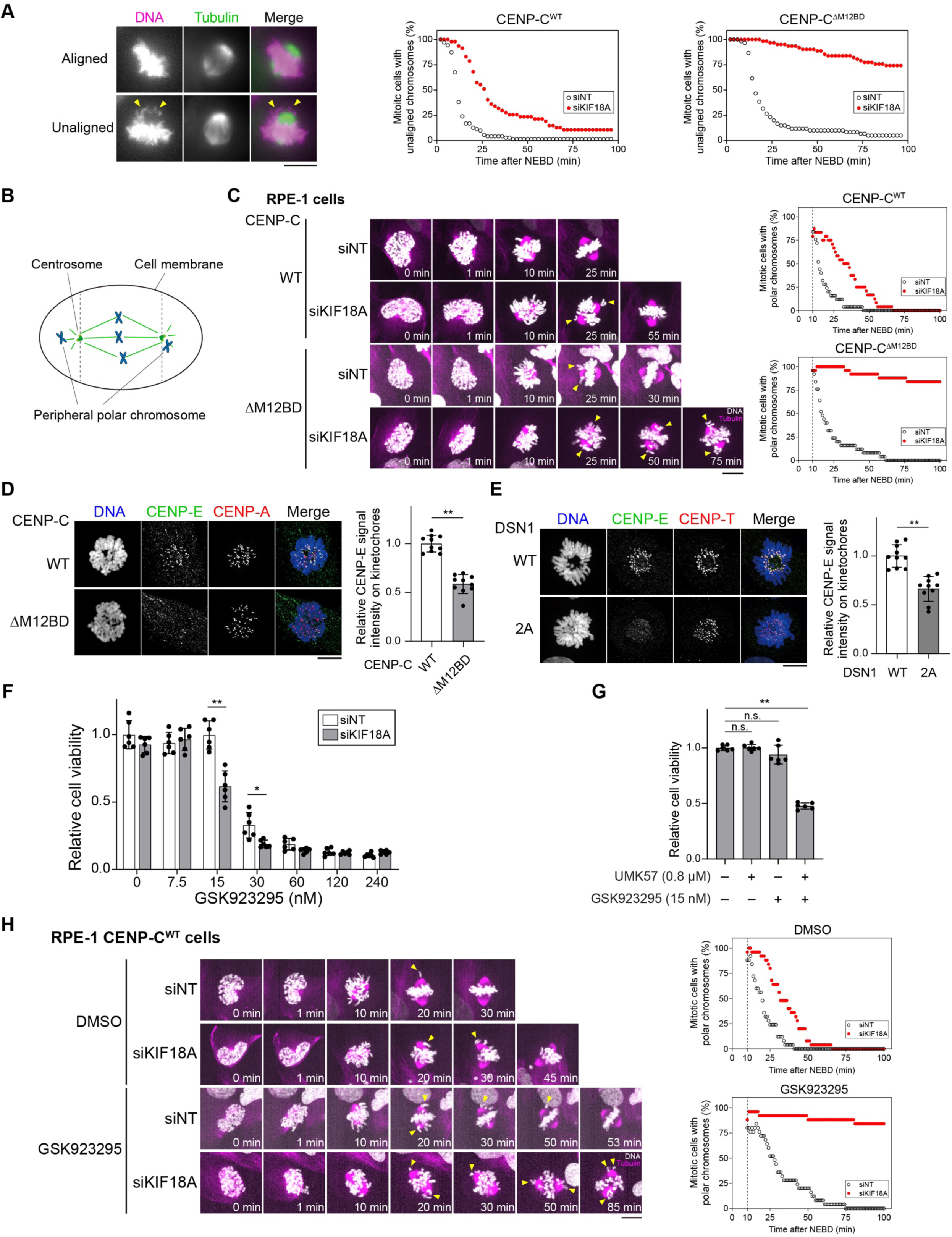
Reduction of CENP-E kinetochore localization is a potential cause of KIF18A-KD sensitivity in RPE-1 CENP-C^ΔM12BD^ cells. (A) Chromosomes alignment in RPE-1 CENP-C^WT^ or CENP-C^ΔM12BD^ cells treated with siNT or siKIF18A. The time-lapse images in Figure 3C were inspected to assess chromosome alignment. The representative time-lapse images of unaligned or aligned chromosomes in RPE-1 cells are shown. Scale bar, 10 μm. The cells with unaligned chromosomes in RPE-1 CENP-C^WT^ or CENP-C^ΔM12BD^ cells treated with siNT or siKIF18A were quantified. (B) Schematic representation of peripheral chromosomes, which are located in the outside zone of two spindle poles (centrosomes). (C) Representative high-resolution time-lapse images to observe peripheral chromosome congression in RPE-1 CENP-C^WT^ or CENP-C^ΔM12BD^ cells treated with siNT or siKIF18A. Arrow heads indicate polar chromosomes. Scale bar, 10 μm. Population of cells with polar chromosomes in RPE-1 CENP-C^WT^ or CENP-C^ΔM12BD^ cells treated with siNT or siKIF18A were quantified (CENP-C^WT^ cells with siNT: n = 25 cells, CENP-C^WT^ cells with siKIF18A: n = 24 cells, CENP-C^ΔM12BD^ cells with siNT: n = 25 cells, CENP-C^ΔM12BD^ cells with siKIF18A: n = 25 cells). (D) CENP-E localization in RPE-1 CENP-C^WT^ or CENP-C^ΔM12BD^ cells. CENP-E was stained with an anti-CENP-E antibody (green). mScarlet-CENP-A was used as a kinetochore marker (CENP-A, red). DNA was stained with DAPI (blue). Scale bar, 10 μm. Signal intensity of CENP-E at mitotic kinetochores was quantified (Mean and SD, two-tailed Student’s t-test, CENP-C^WT^ cells: n = 10 cells, CENP-C^ΔM12BD^ cells: n = 10 cells; \*\**p* < 0.01). (E) CENP-E localization in DSN1^WT^ or DSN1^2A^ RPE-1 cells. CENP-T was used as a kinetochore marker (CENP-T, red). CENP-E (green) and DNA (blue) were stained as in (D). Scale bar, 10 μm. Signal intensity of CENP-E at mitotic kinetochores was quantified (Mean and SD, two-tailed Student’s t-test, DSN1^WT^ cells: n = 10 cells, DSN1^2A^ cells: n = 10 cells; \*\**p* < 0.01). (F) Cell viability of RPE-1 CENP-C^WT^ cells in the presence of GSK923295 at various concentrations after treatment of siNT or siKIF18A (Mean and SD, two-tailed Student’s t-test, each sample size: n = 6; \**p* < 0.1; \*\**p* < 0.01). (G) Cell viability assay of RPE-1 CENP-C^WT^ cells with treatment of 0.8 μM UMK57 and 15 nM GSK923295 (Mean and SD, two-tailed Student’s t-test, RPE-1 cells neither UMK57 nor GSK923295: n = 6, RPE-1 cells with UMK57 but without GSK923295: n = 6, RPE-1 cells without UMK57 but with GSK923295: n = 6, RPE-1 cells with both UMK57 and GSK923295: n = 6; n.s., non-significant; \*\**p* < 0.01). (H) Representative high-resolution time-lapse images to observe peripheral chromosome congression in RPE-1 CENP-C^WT^ cells treated with siNT or siKIF18A in the presence or absence of a CENP-E inhibitor GSK923295. Arrowheads indicate peripheral chromosomes. Scale bar, 10 μm. The population of cells with polar chromosomes in the indicated experimental samples was quantified (CENP-C^WT^ cells with siNT: n = 25 cells, CENP-C^WT^ cells with siKIF18A: n = 25 cells, CENP-C^ΔM12BD^ cells with siNT: n = 25 cells, CENP-C^ΔM12BD^ cells with siKIF18A: n = 25 cells).

To closely evaluate the dynamics of unaligned chromosomes, we also performed high-resolution time-lapse imaging (Figures 5B and 5C). After NEBD, peripheral chromosomes (Figure 5B), which reside near one of the spindle poles, appeared and were aligned to the metaphase plate during mitotic progression in CENP-C^WT^ RPE-1 cells treated with siNT (Figure 5C). The siKIF18A treatment delayed peripheral chromosome congression in RPE-1 CENP-C^WT^ cells, suggesting that KIF18A is involved in the prevention of peripheral chromosome formation and/or peripheral chromosome congression (Figure 5C). The peripheral chromosome congression in RPE-1 CENP-C^ΔM12BD^ cells treated with siNT occurred with a slight delay, compared with that in RPE-1 CENP-C^WT^ cells treated with siNT (Figure 5C). In contrast, the peripheral chromosomes failed to congress to the spindle equator in RPE-1 CENP-C^ΔM12BD^ cells treated with siKIF18A (Figure 5C). The sustained unaligned peripheral chromosomes would cause prolonged mitotic arrest, leading to severe mitotic defects, such as cohesion fatigue (Figure 3G). These observations suggest that the M12BD of CENP-C is required for efficient peripheral chromosome congression and impaired chromosome congression may be a cause of KIF18A-KD sensitivity in RPE-1 CENP-C^ΔM12BD^ cells.

Multiple motor proteins regulate microtubule dynamics and chromosome movements to promote chromosome congression during early prometaphase.^49,50^ CENP-E is a plus-end-directed kinesin that contributes to the conversion of lateral to end-on kinetochore-microtubule attachments, chromosome congression, and the maintenance of alignment at the spindle equator.^32–34,51^ We hypothesized that weakened CENP-E function might be a potential cause of the KIF18A-KD sensitivity in RPE-1 CENP-C^ΔM12BD^ cells. In support of this hypothesis 1) chromosome congression was delayed in RPE-1 CENP-C^ΔM12BD^ cells (Figures 5A and 5C), similar to CENP-E depleted cells^32–34^; 2) a pool of CENP-E is recruited to kinetochores via BubR1, which binds to KNL1,^52,53^ and shows reduced localization in RPE-1 CENP-C^ΔM12BD^ cells^23^; 3) CENP-E localization to kinetochores additionally relies on the RZZ complex^54^, which is also reduced in RPE-1 CENP-C^ΔM12BD^ cells (Figure S5G); 4) chromosome congression mediated by CENP-E depends on stabilized microtubules,^6^ which are diminished in KIF18A-depleted cells.^28,29,45^

To test this hypothesis, we examined CENP-E levels at kinetochores in RPE-1 CENP-C^WT^ and CENP-C^ΔM12BD^ cells. Immunostaining for CENP-E revealed that CENP-E levels at prometaphase kinetochores were significantly reduced in RPE-1 CENP-C^ΔM12BD^ cells (Figures 5D and S7E). In addition, CENP-E levels at kinetochores were also reduced in RPE-1 DSN1^2A^ cells (Figure 5E), which were sensitive to *KIF18A* knockdown (Figure 3F). These results are consistent with the idea that the CENP-E reduction at kinetochores causes the KIF18A-KD sensitivity in RPE-1 CENP-C^ΔM12BD^ cells.

If the CENP-E reduction at kinetochores is the cause of the KIF18A-KD sensitivity in CENP-C^ΔM12BD^ cells, suppression of CENP-E function alone should also lead to KIF18A-KD sensitivity. To test this, we treated RPE-1 CENP-C^WT^ cells with GSK923295, an inhibitor of CENP-E motor activity. Treatment with sublethal doses of GSK923295 reduced cell viability after siKIF18A treatment in RPE-1 CENP-C^WT^ cells (Figure 5F). Thus, CENP-E inhibition results in KIF18A-KD sensitivity in RPE-1 cells, suggesting a functional relationship between CENP-E and KIF18A. This idea was further confirmed by UMK57 treatment, which increased microtubule dynamics. Sublethal-doses of either GSK923295 or UMK57 alone did not reduce cell viability (Figure 5G). However, their simultaneous treatment resulted in a significant reduction in cell viability (Figure 5G).

We also analyzed chromosome dynamics after NEBD in RPE-1 CENP-C^WT^ cells treated with a sublethal dose of GSK923295, using high-resolution time-lapse imaging (Figure 5H). Although the dose of GSK923295 treatment delayed peripheral chromosome congression, eventually the chromosomes aligned to the equator (Figure 5H). By contrast, both siKIF18A and GSK923295 treatment sustained peripheral chromosomes in RPE-1 CENP-C^WT^ cells during the observation (Figure 5H).

Overall, our results suggest that CENP-E and KIF18A work together to facilitate chromosome congression during early prometaphase for proper mitotic progression. Based on these results, we conclude that the impaired CENP-E localization at kinetochores is a cause of the KIF18A-KD sensitivity in RPE-1 CENP-C^ΔM12BD^ cells.

### CENP-E levels were significantly reduced in cancer cell lines sensitive to *KIF18A*-knockdown

The sensitivity to *KIF18A* knockdown or inactivation is observed in multiple cancer cell lines.^41,55,56^ Previous studies found that aneuploidy and weakened anaphase-promoting complex/cyclosome activity were the causes of the vulnerability to KIF18A inactivation in some cancer cell lines.^41,55,56^ Our results indicate that reduced CENP-E kinetochore localization also results in KIF18A-KD sensitivity in RPE-1 cells. This prompted us to assess CENP-E levels at kinetochores in several cancer and non-transformed cell lines. As shown in Figure 6A, in addition to RPE-1 and K562 cells, U2OS, A549, and TIG-3 cells were insensitive to *KIF18A* knockdown. By contrast, HeLa, HT1080, OVCAR-3, HT29, and HCC1806 were sensitive to *KIF18A* knockdown (Figure 6A). The CENP-E levels at kinetochores were significantly lower in the KIF18A-KD sensitive cells, but not in the KIF18A-KD insensitive cells, compared to those in RPE-1 cells (Figure 6B). We also measured CENP-C and CENP-T levels at kinetochores in these cells (Figure 6C and 6D). CENP-C and CENP-T were reduced in HT29 and HCC1806, but not in HeLa, HT1080 and OVCAR-3 cells, suggesting that there were various ways to reduce CENP-E localization at kinetochores among cancer cell lines. In any case, our results demonstrate a correlation between the reduction in kinetochore-localized CENP-E and the KIF18A-KD sensitivity in all tested cancer cell lines. Thus, reduction of CENP-E levels or activity at kinetochores could serve as potential markers for vulnerability to KIF18A inactivation in cancer cell lines.

**Figure 6.**
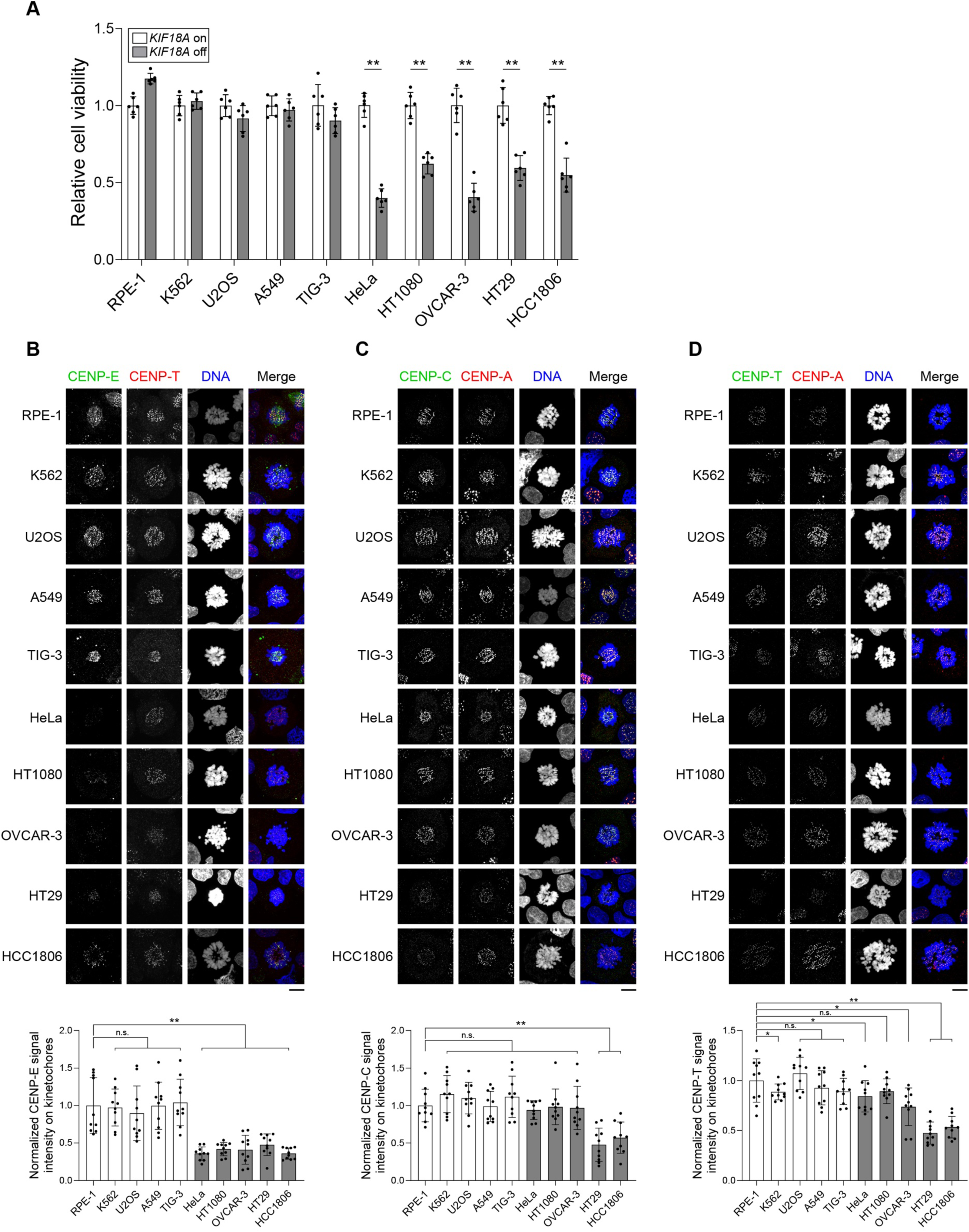
CENP-E levels are reduced in cancer cell lines sensitive to *KIF18A* knockdown. (A) Cell viability of RPE-1, K562, U2OS, A549, TIG-3, HeLa, HT1080, OVCAR-3, HT29, and HCC1806 cells with or without *KIF18A* knockdown. (Mean and SD, two-tailed Student’s t-test, each sample size: n = 6; \*\**p* < 0.01). (B-D) CENP-E (B), CENP-C (C), and CENP-T (D) localization in RPE-1, K562, U2OS, A549, TIG-3, HeLa, HT1080, OVCAR-3, HT29 and HCC1806 cells. The cell lines were treated with 50 μM Monastrol for 2 h to enrich prometaphase cells. Each cell line was cytospan together with Monastrol-treated RPE-1 cells expressing mScarlet-CENP-A as an internal control for immunostaining. CENP-E was stained with an anti-CENP-E antibody (green) (B). CENP-C was stained with an anti-CENP-C antibody (green) (C). CENP-T was stained with an anti-CENP-T antibody (green) (D). DNA was stained with DAPI (blue). CENP-T or CENP-A was stained as a kinetochore marker (red). Scale bar, 10 μm. CENP-E, CENP-C or CENP-T signal intensities at mitotic kinetochores were quantified and normalized with those of the internal control RPE-1 in each sample (Mean and SD, two-tailed Student’s t-test, each cell line: n = 10 cells; n.s., non-significant; \**p* < 0.1; \*\**p* < 0.01).

## Discussion

In this study, we identified genes whose knockout exhibits synthetic mitotic defects with a CENP-C mutant lacking its Mis12 complex binding domain in human cells. Among the identified gene targets, *KIF18A* knockout caused severe lethality in CENP-C^ΔM12BD^ cells. Our in-depth characterization found that reduced levels of the kinesin motor CENP-E in CENP-C^ΔM12BD^ cells caused a mitotic defect when KIF18A was depleted. Based on these findings, we propose that the CENP-C pathway plays a role in proper mitotic progression, with KIF18A and CENP-E acting cooperatively to promote the congression of peripheral chromosomes during early prometaphase (Figure 7).

**Figure 7.**
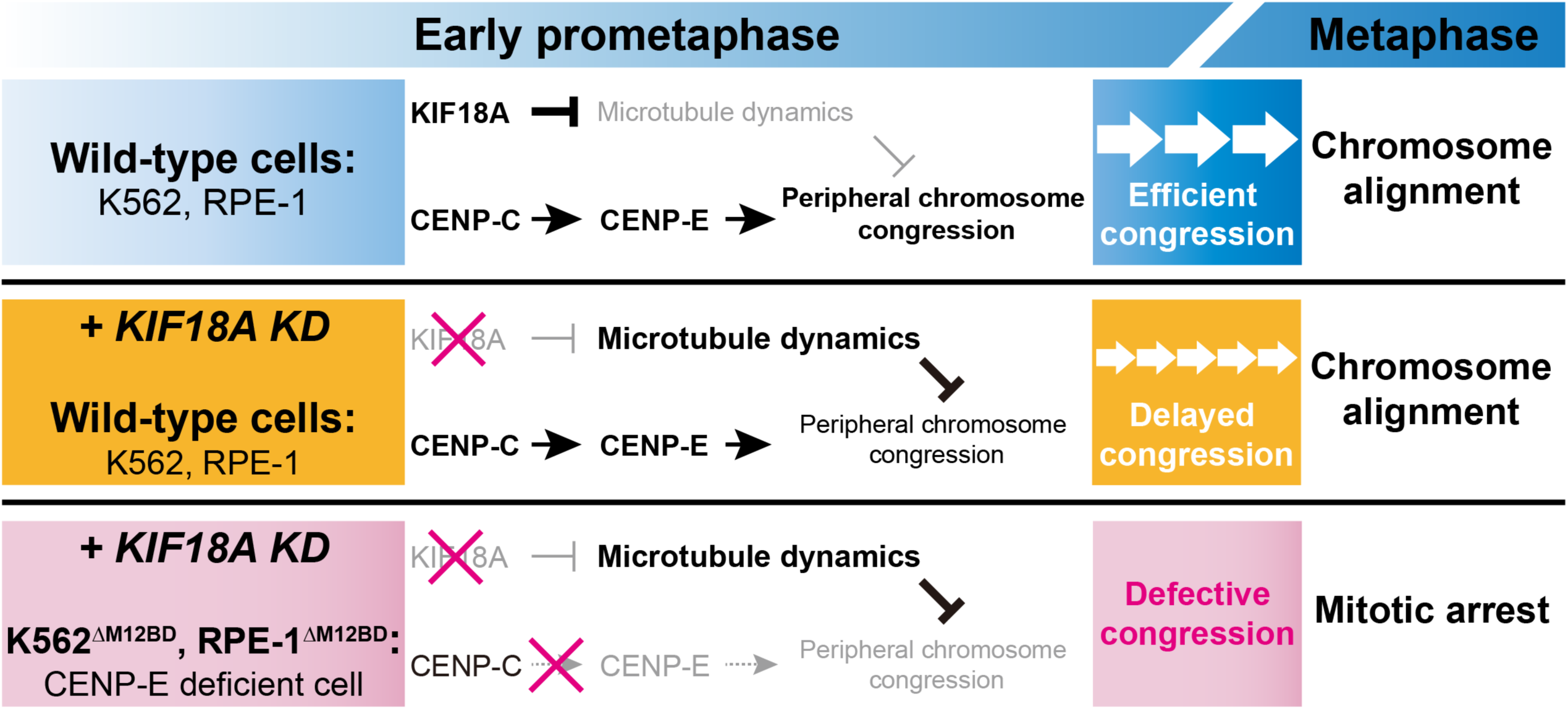
KIF18A and CENP-E cooperatively function for the congression of peripheral chromosomes at early prometaphase. In wild-type K562 and RPE-1 cells (Middle), although KIF18A depletion enhances microtubule dynamics and increases peripheral chromosomes, CENP-E downstream of CENP-C compensates for KIF18A depletion, supporting chromosome congression and alignment, albeit with a delay in the congression. By contrast, in CENP-E deficient cells (Bottom: K562 CENP-C^ΔM12BD^ and RPE-1 CENP-C^ΔM12BD^ cells), KIF18A depletion disrupts chromosome congression due to weakened CENP-E activity downstream of CENP-C at kinetochores, leading to mitotic arrest and subsequent cell death. Based on these observations, we propose that the CENP-C pathway plays a role in chromosome congression, with KIF18A and CENP-E acting cooperatively to promote the congression of peripheral chromosomes during early prometaphase in wild-type K562 and RPE-1 cells (Upper). KIF18A suppresses microtubule dynamics to facilitate congression and/or prevent the formation of peripheral chromosomes. CENP-E also plays a key role in efficient chromosome congression by gliding kinetochores along microtubules.

For the CRISPR screen, we took advantage of the unique characteristics of CENP-C^ΔM12BD^. Although the deletion of the CENP-C M12BD reduced the recruitment of outer kinetochore complex KMN and caused minor mitotic defects, it did not affect overall cell growth in chicken cells, human cells, or mice in vivo.^21–23^ This domain is dispensable because another CCAN protein, CENP-T, also recruits KMN to kinetochores. KMN bound to CENP-T is necessary and sufficient for cell viability, whereas the KMN bound to CENP-C is dispensable.^21,22^ However, a recent detailed analysis of CENP-C M12BD in human cells revealed that the domain facilitated chromosome alignment through Aurora B kinase.^23^ In current our CRISPR screen, in addition to *KIF18A*, we also identified several genes involved in spindle formation and chromosome congression (*KIF15*, *NEDD1*, *HAUS8*, *NDE1*, *KNTC1*, *ZW10*).^57–67^ This suggests that CENP-C contributes to chromosome congression during prometaphase to facilitate bioriented end-on attachment mediated through CENP-T.

KMN recruitment to CENP-C M12BD is promoted by Aurora B phosphorylation of DSN1, a subunit of Mis12C.^25–27,42,43^ In RPE-1 cells, Aurora B protein and phosphorylated DSN1 are enriched at kinetochores on unaligned chromosomes compared with aligned chromosomes.^68^ This suggests a dynamic modulation of KMN recruitment to CENP-C during chromosome congression in early prometaphase. Interestingly, an enrichment of Aurora B and phosphorylated DSN1 at unaligned kinetochores was not found in cancer cell lines, including HeLa cells.^68^ As the dynamic KMN modulation via CENP-C does not happen in RPE-1 CENP-C^ΔM12BD^ cells, these cells could provide a cancer model to study chromosome instability.

The kinetochore localization of many proteins, including KMN and Aurora B, is reduced in RPE-1 CENP-C^ΔM12BD^ cells. As Aurora B and phosphorylated DSN1 are both enriched at kinetochores on unaligned chromosomes,^68^ their recruitment via CENP-C might be important for chromosome congression. Interestingly, an enrichment of Aurora B and phosphorylated DSN1 at unaligned kinetochores was not found in cancer cell lines, including HeLa cells. This observation is similar to that of RPE-1 CENP-C^ΔM12BD^ cells. Indeed, *KIF18A* knockdown or inhibition kills chromosomally unstable cancer cells,^55,56,69^ and was also identified as a gene whose knockout causes synthetic lethality with CENP-C^ΔM12BD^ in our CRISPR screen.

KIF18A is a Kinesin-8 family plus-end-directed motor protein that controls microtubule dynamics during mitosis.^28–31^ Studies with KIF18A depletion demonstrated that KIF18A stabilizes kinetochore-microtubule attachments and suppresses the pre-anaphase oscillation of end-on-attached chromosomes to facilitate chromosome alignment.^28,29^ Our genome-wide screening in the CENP-C mutant cells and our detailed downstream analysis revealed that KIF18A functions in parallel with the plus-end-directed kinesin CENP-E (Kinesin-7).^32,70,71^ CENP-E plays a major role in the congression of unaligned chromosomes, which remain near the spindle pole following NEBD. CENP-E promotes chromosome alignment through its motor activity-dependent kinetochore-gliding along spindle microtubules in early prometaphase.^32–34^ This suggests that, in addition to its established role in chromosome oscillations in metaphase, KIF18A plays a role in congression of peripheral chromosomes or the prevention of peripheral chromosome formation during the early prometaphase in cooperation with CENP-E. Consistent with this idea, we observed increased unaligned chromosomes and polar chromosomes after NEBD in KIF18A knockdown cells (Figure 5A and 5C). However, the role of KIF18A in the alignment of peripheral chromosomes has been missed due to the overlapping roles of CENP-E, which appears to compensate for KIF18A depletion in early prometaphase.

Our previous study revealed that Aurora B levels were reduced at centromeres in CENP-C^ΔM12BD^ cells.^23^ We tried to examine whether the Aurora B reduction was a cause of the KIF18A-KD sensitivity. However, even low doses of AZD1152, an Aurora B inhibitor, affected cell viability in the control siNT-treated cells; we could not assess the effect of reduced Aurora B activity in siKIF18A-treated RPE-1 CENP-C^WT^ cells (Figure S7D). Since Aurora kinase activity has been shown to release CENP-E from its autoinhibition,^72^ it is possible that the Aurora B reduction could be the primary cause of the KIF18A-KD sensitivity in CENP-C^ΔM12BD^ cells. A recent study proposed that the CENP-E function was opposed by Aurora kinase activity.^73^ These observations imply that Aurora B could regulate CENP-E both positively and negatively, complicating the interpretation of the effect of Aurora B inhibition on the KIF18A-KD sensitivity. Although whether and how Aurora B is related to the KIF18A-KD sensitivity remain to be clarified, our results clearly demonstrate the reduced CENP-E localization as a cause of the KIF18A-KD sensitivity in CENP-C^ΔM12BD^ cells.

How does KIF18A contribute to chromosome congression and alignment in early prometaphase in cooperation with CENP-E? After NEBD, most chromosomes are located between two spindle poles where they rapidly establish end-on attachments for biorientation. However, a subset of chromosomes is left behind closer to one of the poles.^74,75^ KIF18A may promote the establishment of end-on attachments by stabilizing kinetochore-microtubule interaction, leading to direct chromosome congression to the spindle equator. This KIF18A function at early prometaphase may assist CENP-E to glide the peripheral chromosomes along the stable K-fibers to promote their congression. A recent study proposed that the critical function of CENP-E in chromosome congression was the formation of end-on microtubule attachments rather than gliding the chromosome through lateral attachments.^73^ In this case, KIF18A could support CENP-E function by stabilizing kinetochore-microtubule attachments. Alternatively, KIF18A may regulate CENP-E protein stability. A previous study reported that KIF18A depletion reduced CENP-E protein levels and its localization at mitotic kinetochores.^76^ *KIF18A* knockdown may further reduce CENP-E levels in CENP-C^ΔM12BD^ RPE-1 cells, leading to mitotic defects. In either case, KIF18A has a genetic interaction with CENP-E and contributes to peripheral chromosome congression.

Since several cancer cell lines, but not non-transformed cell lines, are sensitive to *KIF18A* knockdown or small KIF18A small-chemical inhibitors,^55,56,69,77,78^ KIF18A has been proposed as a potential therapeutic target in cancers. However, not all cancer cell lines are sensitive to KIF18A inhibition. Therefore, understanding the molecular mechanisms for the vulnerability of KIF18A inhibition in cancer cell lines is critical for effective cancer therapy targeting KIF18A. A recent study demonstrated that weakened anaphase-promoting complex/cyclosome and/or persistent SAC activation resulted in the prolonged mitosis leading to the vulnerability of KIF18A inhibition.^41^ Indeed, preventing SAC activation suppressed sensitivity to KIF18A knockdown in our RPE-1 CENP-C^ΔM12BD^ cells (Figure 3I). In RPE-1 CENP-C^ΔM12BD^ cells, *KIF18A* knockdown increased unaligned chromosomes, resulting in SAC activation and a prolonged mitosis. In addition, we demonstrated that at least five cancer cell lines that are vulnerable to *KIF18A* knockdown also exhibited reduced CENP-E levels at kinetochores, consistent with a recent report.^79^ Our findings suggest that the reduction of CENP-E levels at mitotic kinetochores is one of the determinants of KIF18A dependency in certain cancer cell lines, nominating CENP-E as a potential biomarker for effective cancer therapy with KIF18A inhibition.

Since chromosome segregation is ensured by complex and redundant mechanisms, the identification of factors involved in the mechanisms and understanding their relationship is challenging. Our CRISPR screen with CENP-C^ΔM12BD^ cells reveals the hidden interplay regulating chromosome congression. Thus, our approach using a mutant with modest mitotic defects provides a powerful tool to reveal new insights for understanding the complicated mechanisms that underlie chromosome segregation.

## Limitation of the Study

Although we proposed that KIF18A and CENP-E cooperatively function for congression of peripheral chromosomes in early prometaphase, understanding of detailed molecular mechanisms to achieve chromosome congression is still limited. Whereas we proposed a few possibilities of how KIF18A and CENP-E function cooperatively, clarifying these possibilities would be the next key questions to be addressed.

## Resource Availability

### Lead contact

Requests for further information should be directed to the lead contact, Tatsuo Fukagawa.

## Materials Availability

All materials generated in this study are available from the lead contact upon request.

## Data and Code Availability

No new code was generated in this study.

Any additional information required to reanalyze the data reported in this paper is available from the lead contact upon request.

## Acknowledgments

The authors are very grateful to members of the Fukagawa Lab for the fruitful discussion. We also thank R. Fukuoka and Y. Kubota for their technical assistance and T. Hirota and K. Tanaka for providing cell lines. This work was supported by CREST of JST (JPMJCR21E6), JSPS KAKENHI Grant Numbers 20H05389, 21H05752, 22H00408, 22H04692, and 24H02281 to TF, JSPS KAKENHI Grant Numbers 16K18491, 21H02461 22H04672, and 24K02005, Takeda Science Foundation and Daiichi Sankyo Foundation of Life Science to MH, and NIH/NIGMS (R35GM126930) to IMC.

## Author contributions

Conceptualization: M.H., T.F.; Investigation: J.M., K.-C.S., H.R.K., W.K., Y.T.; Formal analysis: J.M., M.H., K.-C.S., H.R.K.; Resources: J.M., W.K., Y.T.; Data curation: J.M., M.H.; Writing - original draft: M.H., T.F.; Writing - review & editing: M.H., J.M., T.F., I.M.C.; Supervision: M.H., T.F.; Project administration: T.F.; Funding acquisition: M.H., T.F. I.M.C.

## Star Methods

### EXPERIMENTAL MODEL AND STUDY PARTICIPANT DETAILS

#### Cell culture

Human K562 cells were maintained in a culture medium containing RPMI-1640 (Nacalai Tesque) supplemented with 10% fetal bovine serum (Sigma-Aldrich) and Penicillin (100 unit/ml)-Streptomycin (100 μg/ml) (Thermo Fisher Scientific) and cultured at 37°C with 5% CO_2_. Human hTERT-RPE-1, Lenti-X 293T (Takara Bio), HeLa Kyoto, HT1080, U2OS, TIG-3 and A549 cell lines were maintained in a culture medium containing DMEM (Nacalai Tesque) supplemented with 10% fetal bovine serum (Sigma-Aldrich) and Penicillin (100 unit/ml)-Streptomycin (100 μg/ml) (Thermo Fisher Scientific) and cultured at 37°C with 5% CO_2_. For the large-scale lentiviral preparation, Lenti-X 293T cells were maintained in a culture medium containing IMDM (Thermo Fisher Scientific) supplemented with 20% fetal bovine serum (Sigma-Aldrich), and cultured at 37°C with 5% CO_2_. Human OVCAR-3, HT29 and HCC1806 cell lines were maintained in a culture medium containing RPMI1640 (Nacalai Tesque) supplemented with 10% fetal bovine serum, and cultured at 37°C with 5% CO_2_.

#### Plasmid constructions for cell transfection

To establish K562 cells expressing CENP-C^ΔM12BD^, CRISPR/Cas9-mediated gene editing was utilized by employing two independent pX330 (a gift from Feng Zhang, Addgene plasmid, #42230),^80^ one pX330 containing single-guide RNA (sgRNA) targeting a genomic sequence in the *CENP-C* exon2 (pX330_sgCCExon2), another pX330 containing sgRNA targeting a genomic sequence in the *CENP-C* exon 4 (pX330_sgCCExon4). sgRNA for *CENP-C* genome editing was designed using CRISPOR.^81^

To establish an inducible Cas9 mediated gene knockout system in K562 cell, a Doxycycline-inducible Cas9 cassette and a landing pad with *attB* sequences for integration of sgRNA sequences were cloned into the human *AAVS1* backbone plasmid using In-Fusion HD Cloning Kit (Takara Bio). The Doxycycline-inducible Cas9 cassette contains *Cas9* with the Tet-responsible (Tet-On) promoter followed by *rtTA-T2A*-neomycin resistance genes (*NeoR*) with the *EF1a* promoter. The cassette was amplified from HP138-neo (Addgene plasmid, #134247)^82^ by PCR. The landing pad contains *HSV TK* with CMV promoter flanked with two *attB* sequences. The Doxycycline-inducible Cas9 cassette and landing pad were cloned into the human *AAVS1* backbone plasmid which was derived from digested pAAVS1-NDi-CRISPRi (Gen1) (a gift from Bruce Conklin, Addgene plasmid, #73497)^83^ by the restriction enzyme EcoRV and HindIII by using In-Fusion HD Cloning Kit (Takara Bio) (pAAVS1_Tet-Cas9). This construct was integrated into the *AAVS1* locus by CRISPR/Cas9-mediated homologous recombination, employing pX330 containing sgRNA targeting a sequence around the *AAVS1* exon1 (pX330_sgAAVS1).

To generate a donor plasmid congaing *attP* and three sgRNA sequences targeting a gene, each sgRNA was cloned into pX330 (a gift from Feng Zhang, Addgene plasmid, #42230)^80^ independently. Three sgRNAs were designed for a gene at different exons using CRISPOR.^81^. Using each pX330 with sgRNA as a template, the sequence with the U6 promoter and sgRNA was amplified with PCR. The amplified sequences (three for each gene target) were fused and flanked with two *attP* sequences by using In-Fusion HD Cloning Kit (Takara Bio) (pattP_tri-gRNA).

To express mScarlet-fused human full-length CENP-A under the control of the endogenous CENP-A promoter in human RPE-1 cells, the sequence of *mScarlet-fused CENP-A* cDNA followed by puromycin resistance gene (*PuroR*) or *NeoR* expression cassette driven by the *beta-actin* promoter (*ACTB*) was flanked with 5’ and 3’ *CENP-A* homology arm fragments (approximately 1 kb each) which were surrounding the *CENP-A* start codon was cloned into the pBluescript II SK (pBSK) by using In-Fusion HD Cloning Kit (Takara Bio) (pBSK_mScarlet-CENP-A).^23^ The construct was integrated into the endogenous *CENP-A* genome locus by CRISPR/Cas9-mediated homologous recombination with pX330 containing sgRNA targeting a sequence around the *CENP-A* start codon (pX330_sgCENP-A).^23^

To express mScarlet-fused human full-length DSN1 (DSN1^WT^) or S100A/S109A (DSN1^2A^) mutant under the control of the endogenous *DSN1* promoter in RPE-1 cells, the cDNA of *mScarlet-fused DSN1 WT* or *2A* mutant followed by *PuroR* or *NeoR* expression cassette driven by the *ACTB* promoter was flanked with 5’ and 3’ homology arm fragments (approximately 1 kb each) which is surrounding the *DSN1* start codon was cloned into the pBSK (pBSK_mScarlet-DSN1^WT^ or pBSK_mScarlet-DSN1^2A^) by using In-Fusion HD Cloning Kit (Takara Bio). The constructs were integrated into the endogenous *DSN1* gene locus by CRISPR/Cas9-mediated homologous recombination with pX330 containing sgRNA targeting a sequence around the *DSN1* start codon (pX330_sgDSN1).^23^

For KIF18A analysis with siRNA knockdown, siRNA targeting sequences of *EGFP-fused full-length KIF18A* cDNA were modified with synonymous mutations to make siRNA-resistant KIF18A construct (KIF18A^WT^). The synonymous mutations were designed by Synonymous Mutation Generator.^84^ To express EGFP-fused KIF18A^WT^ under the control of CMV promoter in RPE1 cells, the *EGFP-fused KIF18A^WT^*cDNA followed by the Blasticidin S resistance gene (*BsR*) expression cassette driven by the *ACTB* promoter flanked with 5’ and 3’ homology arm fragments of *AAVS1* locus (approximately 1 kb each) was cloned into the pBSK by using In-Fusion HD Cloning Kit (Takara Bio) (pBSK_CMV-KIF18A^WT^-BsR). To express EGFP-fused human KIF18A **Δ**1-362 (KIF18A**^Δ^**^1-362^), **Δ**PP1 (KIF18A**^Δ^**^PP1^), S674A/S684A (KIF18A^2A^) or **Δ**PP1 + S674A/S684A (KIF18A**^Δ^**^PP1+2A^) mutant, each siRNA-resistant mutant construct was cloned as EGFP-fused KIF18A^WT^. (pBSK_CMV-KIF18A**^Δ^**^1-362^-BsR, pBSK_CMV-KIF18A**^Δ^**^PP1^-BsR, pBSK_CMV-KIF18A^2A^-BsR and pBSK_CMV-KIF18A**^Δ^**^PP1+2A^-BsR). Each construct was integrated into the endogenous *AAVS1* gene locus by CRISPR/Cas9-mediated homologous recombination with (pX330_sgAAVS1).

To clone an sgRNA sequence to pX330 (a gift from Feng Zhang, Addgene plasmid, #42230),^80^ forward and reverse oligos including sgRNA sequence and flanking sequences^80^ were incubated with T4 Polynucleotide Kinase (TOYOBO) at 37°C for 30 min, then annealed under the following sequential protocol: 95°C for 5 min, 75°C for 5 min, 55°C for 5 min, finally 35°C for 5 min. Annealed DNAs were ligated with pX330 (a gift from Feng Zhang, Addgene plasmid, #42230),^80^ which is digested by BpiI (Thermo Fisher Scientific). All sgRNA sequences and oligo sequences used in this study are listed in Table S2.

### METHOD DETAILS

#### Generation of cell lines

To establish K562 cells expressing CENP-C**^Δ^**^M12BD^, wild-type K562 cells (K562 WT cells) were co-transfected with pX330_sgExon2 and pX330_sgExon4 using Neon Transfection System (Thermo Fisher Scientific) with 1 pulse (1000 V, 50 msec), as previously described.^22^ Single-cell clones were isolated from transfected cells (K562 CENP-C**^Δ^**^M12BD^ cells).

To establish a Doxycycline-inducible Cas9-mediated gene knockout system in K562 cells, K562 WT or K562 CENP-C**^Δ^**^M12BD^ cells were co-transfected with pX330_sgAAVS1 and pAAVS1_Tet-Cas9 using Neon Transfection System (Thermo Fisher Scientific) with 1 pulse (1000 V, 50 msec). Transfected cells were selected in a medium containing 1 mg/ml G418 (Santa Cruz Biotechnology) to isolate single-cell clones (K562 WT/Cas9 cells, K562 CENP-C**^Δ^**^M12BD^/Cas9 cells). To integrate sgRNA sequences to the *attB* landing pad in K562 WT/Cas9 or CENP-C**^Δ^**^M12BD^/Cas9 cells, cells were co-transfected with pattP_tri-gRNA for each target gene and Bxb1 expression plasmid pCAG-NLS-HA-Bxb1 (a gift from Pawel Pelczar, Addgene plasmid, #51271)^85^ using Neon Transfection System (Thermo Fisher Scientific) with 1 pulse (1000 V, 50 msec). Transfected cells were selected in a medium containing 500 nM FIAU (Sigma-Aldrich) to isolate single-cell clones.

To establish RPE-1 cell lines expressing mScarlet-fused CENP-A under the control of the endogenous *CENP-A* promoter, the RPE-1 cells were co-transfected with pX330_sgCENP-A and two pBSK_mScarlet-CENP-A plasmids, each containing *PuroR* or *NeoR*, using Neon Transfection System (Thermo Fisher Scientific) with 6 pulses (1400 V, 5 msec). Transfected cells were then selected in a medium containing 2 mg/ml puromycin (Takara Bio) and 500 μg/ml G418 (Santa Cruz Biotechnology) to isolate single-cell clones (RPE-1 mScarlet-CENP-A cells).^23^

To establish RPE-1 expressing either CENP-C^WT^ or CENP-C^ΔM12BD^ under the control of the endogenous *CENP-C* promoter, and also expressing mScarlet-CENP-A, RPE-1 cells expressing mScarlet-CENP-A under the control of the endogenous *CENP-A* promoter (RPE-1 mScarlet-CENP-A cells)^23^ were co-transfected with and pX330_sgCENP-C and two pBSK_CENP-C^WT^ plasmids, each containing *ZeoR* or *HisD,* or two pBSK_CENP-C^ΔM12BD^ under plasmids, each containing *ZeoR* or *HisD,* using Neon Transfection System (Thermo Fisher Scientific) with 6 pulses (1400 V, 5 msec). Transfected cells were selected in a medium containing 10 ng/ml zeocin (Thermo Fisher Scientific) and 1.5 mg/ml L-Histidinol dihydrochloride (Sigma-Aldrich) to isolate single-cell clones (RPE1 mScarlet-CENP-A_CENP-C^WT^ or mScarlet-CENP-A_CENP-C^ΔM12BD^ cells).

To establish RPE-1 cells expressing mScarlet-fused DSN1^WT^ or DSN1^2A^ under the control of the endogenous *DSN1* promoter, RPE-1 cells were co-transfected with pX330_sgDSN1 and two pBSK_mScarlet-DSN1^WT^ plasmids, each containing *PuroR* or *NeoR*, or two pBSK_mScarlet-DSN1^2A^ plasmids, each containing *PuroR* or *NeoR*, using Neon Transfection System (Thermo Fisher Scientific) with 6 pulses (1400 V, 5 msec). Transfected cells were selected in a medium containing 5 μg/ml puromycin (Takara Bio) and 2 mg/ml G418 (Santa Cruz Biotechnology) to isolate single-cell clones (RPE-1 mScarlet-DSN1^WT^ or mScarlet-DSN1^2A^ cells).

To establish RPE-1 cells expressing EGFP-fused KIF18A^WT^, KIF18A^Δ1-362^, KIF18A^ΔPP1^, KIF18A^2A^ or KIF18A^ΔPP1+2A^, RPE-1 cells were co-transfected with pX330_sgAAVS1 and pBSK_CMV-KIF18A^WT^-BsR, pBSK_CMV-KIF18A^Δ1-362^-BsR, pBSK_CMV-Kif18A^ΔPP1^-BsR, pBSK_CMV-Kif18A^2A^-BsR, or pBSK_CMV-KIF18A^ΔPP1+2A^-BsR using Neon Transfection System (Thermo Fisher Scientific) with 6 pulses (1400 V, 5 msec). Transfected cells were selected in a medium containing 20 μg/ml Blasticidin S hydrochloride (Wako) to isolate single-cell clones (RPE-1 EGFP-KIF18A^WT^, EGFP-KIF18A^Δ1-362^, EGFP-KIF18A^ΔPP1^, EGFP-KIF18A^2A^ and EGFP-KIF18A^ΔPP1+2A^ cells).

#### Genotyping

To isolate genomic DNA from K562 or RPE-1 cells, cells were collected and washed with cold PBS. Then cells were centrifuged. The cell pellet was resuspended in 50 mM NaOH and incubated for 10 min at 95°C followed by mixing with a 10% sample volume of 1 M Tris-HCl pH8.0. After centrifugation, supernatant was applied for genome PCR using primers listed in Table S3.

#### Southern blot

1×10^7^ K562 cells were collected and washed with cold PBS, then suspended with lysis buffer (100 mM NaCl, 10 mM Tris-HCl pH 8.0, 25 mM EDTA, 0.5% SDS, 0.1 mg/ml, proteinase K (Sigma-Aldrich)). The lysate was incubated overnight at 50°C with shaking. An equal volume of phenol/chloroform/isoamyl alcohol (25:24:1) (NIPPON GENE) was added and mixed well. The sample was centrifuged for 10 min at 1700*g* at RT. Then, the aqueous phase was isolated and mixed with 1/10 volume of 3 M ammonium acetate and twice the volume of 100% ethanol. Centrifugation at 1700*g* for 2 min was applied to precipitate genomic DNA. The DNA pellet was rinsed with 70% ethanol twice and suspended in TE buffer (10 mM Tris-HCl, 1 mM EDTA, pH 8.0). Dissolved genomic DNA was digested by EcoRI overnight at 37°C. Then, digested DNA was run on 0.8% agarose gel and transferred to nylon membranes (Biodyne B, Cytiva). The membrane was incubated with the DNA probe, which was labeled with alkaline phosphatase with AlkPhos Direct Labeling kit (Cytiva, RPN3680). To detect DNA fragments hybridized with the probe, the membrane was incubated with CDP-Star detection reagent (Cytiva), and signals were visualized using a ChemiDoc Touch imaging system (Bio-Rad). Images were processed using Image Lab (Bio-Rad) and Fiji (v1.53).^86^

#### Cell counting

To count the number of K562 cells, living K562 cells were collected and mixed with an equal volume of 0.4% (w/v) Trypan Blue Solution (Wako). The cell numbers were counted using the Countess II (Thermo Fisher Scientific). Relative cell numbers were calculated based on the cell numbers at time 0 h in each experiment.

#### Immunoblotting

5×10^5^ K562 or RPE-1 cells were collected, washed with cold PBS, and then suspended in 1 x Laemmli Sample Buffer (LSB: 62.5 mM Tris-HCl pH6.8, 10% glycerol, 2% SDS, 5% 2-mercaptoethanol, bromophenol blue) at a final concentration of 1x10^4^ cells/µl. The lysate was sonicated and heated for 5 min at 96°C. The samples were separated by SDS-PAGE using 5-20% gradient polyacrylamide gels (SuperSepAce 5-20%, Wako) and transferred onto a PVDF membrane (Immobilon-P, Merck). After washing the membrane in TBST (50 mM Tris-HCl pH7.5, 150 mM NaCl, 0.1% Tween 20) for 15 min, the membrane was incubated with a primary antibody overnight at 4°C. Following a 15-min wash with TBST, the membrane was incubated with a secondary antibody for 1 h at RT. With a 15-min wash with TBST, the membrane was incubated with ECL Prime Western Blotting Detection Reagent (Cytiva) for 5 min. The signal was detected and visualized using a ChemiDoc Touch imaging system (Bio-Rad) and the image was processed using Image Lab (Bio-Rad) and Fiji (v1.53).^86^

As CENP-C is highly phosphorylated, the CENP-C signals were smeared and unclear on immunoblots. To detect clear CENP-C signals, proteins in cell lysates were dephosphorylated with Lambda Protein Phosphatase (New England Biolabs). Collected cells were suspended in lysis buffer (50 mM NaH_2_PO_4_, 50 mM Na_2_HPO_4_, 300 mM NaCl, 0.1% NP-40, 5 mM 2-mercaptoethanol, 1 x cOmplete EDTA-free proteinase inhibitor (Roche)) at a final concentration of 1x10^5^ cells/μl. The lysate was sonicated, and the supernatant was recovered after centrifugation. Three μl of the supernatant sample was mixed with 20 μl H_2_O, then 3 μl of 10 x NEBuffer and 3 μl of 10 mM MnCl_2_ were added. Then, 1 μl of Lambda Protein Phosphatase was added to the premixed supernatant sample and incubated for 10 min at 30°C according to the manufacturer’s instructions. After mixing with LSB, and heated for 5 min at 96°C. The proteins were detected as described above.

The primary antibodies used in this study were, Guinea pig anti-human CENP-C (1:10,000)^87^, rabbit anti-Cas9 (1:2,000)(Takara Bio), mouse anti-human CENP-A (1:1,000)^87^, rabbit anti-human DSN1 (1:5,000),^88^ rabbit anti-human KIF18A (1:2000) (Bethyl), rabbit anti-human KIF15 (1:2000) (Proteintech), rabbit anti-human NDE1 (1:2000) (Proteintech), rabbit anti-human NEDD1 (1:2000) (Proteintech), rabbit anti-human SPAG5 (1:2000) (Proteintech), rabbit anti-human KNTC1 (1:2000) (Proteintech), rabbit anti-human ASPM (1:2000) (Proteintech), rabbit anti-human TUBGCP6 (1:2000) (Novus biologicals), mouse anti-human NPM1 (1:2000) (Proteintech) and mouse anti-α-tubulin (1:5,000) (Sigma-Aldrich). The secondary antibodies were horseradish peroxidase (HRP) -conjugated goat anti-rabbit IgG (1:5,000) (Jackson ImmunoResearch), HRP-conjugated anti-guinea pig IgG (1:5,000) (Jackson ImmunoResearch) and HRP-conjugated anti-mouse IgG (1:5,000) (Jackson ImmunoResearch). All antibodies were diluted in Signal Enhancer Hikari (Nacalai Tesque) to enhance signal sensitivity and specificity.

#### Immunofluorescence and image acquisition

To localize CENP-C, CENP-T, CENP-A, DSN1, KNL1, HEC1, and MAD2 in K562 cells, K562 cells were cytospan onto slide glasses (MATSUNAMI, S2111) or coverslips (MATSUNAMI, C024361) and then fixed with 3% paraformaldehyde (PFA; Nacalai Tesque) for 10 min at RT, and rinsed with PBS. The cells were permeabilized with 0.5% NP-40/PBS for 10 min at RT, and rinsed with PBS. After blocking with 0.5% BSA/PBS for 5 min at RT, the cells were incubated with primary antibodies diluted in 0.5% BSA/PBS for 1 h at 37°C. The cells were washed three times with 0.5% BSA/PBS, incubated with secondary antibodies diluted in 0.5% BSA/PBS for 1 h at 37°C, and washed three times with 0.5% BSA/PBS. The cells were post-fixed in 3% PFA/PBS for 10 min. DNA was stained with 100 ng/ml DAPI/PBS for 20 min at RT. The stained cells were mounted with VECTASHIELD Mounting Medium (Vector Laboratories).

To localize CENP-C, CENP-T, BubR1, DSN1, KNL1, and HEC1 in RPE-1 cells, RPE-1 cells were seeded onto 35 mm glass-bottom culture dishes (MatTek) and incubated for 24 to 36 h. The cells were fixed with 4% PFA (Electron Microscopy Sciences) in PHEM buffer (60 mM PIPES, 25 mM HEPES, 10 mM EGTA, 2 mM MgCl_2_, pH 6.8) for 10 min at RT. The cells were permeabilized with 0.5% Triton X-100/PBS for 10 min at RT, followed by blocking with blocking solution (3% BSA, 0.1% Triton X-100, 0.1% sodium azide in TBS (50 mM Tris-HCl pH 7.5, 150 mM NaCl)) for 5 to 10 min at RT. The samples were then incubated with primary antibodies in blocking solution overnight at 4°C. After washing the samples with PBST (0.1% Triton X-100/PBS) three times for 5 min each, they were incubated with secondary antibodies in blocking solution for 1 h at RT. Subsequently, the samples were washed in PBST three times for 5 min each, and DNA was stained with 100 ng/ml DAPI/PBS for 10 min at RT. After washing with PBS, cells were mounted with VECTASHIELD Mounting Medium (Vector Laboratories).

To localize ZW10 in RPE-1 cells, the cells were seeded onto 35 mm glass-bottom culture dishes (MatTek) and incubated for 24 h. Then cells were permeabilized for 1 min with warm 0.2% Triton X-100/PHEM buffer, and fixed with 4% PFA/PBS for 10 min. After 3-time wash with PBS, the cells were blocked with 3% BSA/PBS for 1 h at RT. The cells were incubated with primary antibodies diluted in 3% BSA/PBS for 16 h at 4 °C. Subsequently, the cells were washed three times for 5 min each with PBST and incubated with secondary antibodies in 3% BSA/PBS for 1 h at RT. After washing with PBST three times for 5 min each, DNA was stained with 100 ng/ml DAPI/PBS for 10 min at RT. After washing with PBS, cells were mounted with VECTASHIELD Mounting Medium (Vector Laboratories).

We used immunofluorescence to localize EGFP-fused KIF18A proteins in RPE-1 because their EGFP signals were weak. The cells were seeded onto 35 mm glass-bottom culture dishes (MatTek), treated with siKIF18A, and then incubated for 48 h. The cells were fixed in −20°C methanol with 1% PFA for 10 min on ice and washed three times for 5 min each in TBS. The cells were blocked with 20% goat serum (Jackson ImmunoResearch) in antibody-diluting buffer (1% BSA, 0.1% Triton X-100, and 0.1% sodium azide in TBS) for 1 h at RT and incubated with primary antibodies diluted in antibody-diluting buffer for 1 h at RT. Subsequently, the cells were washed with TBS twice for 5 min each and incubated with secondary antibodies for 1 h at RT. After three washs with TBS for 5 min each, DNA was stained with 100 ng/ml DAPI/PBS for 10 min at RT. After washing with PBS, cells were mounted with VECTASHIELD Mounting Medium (Vector Laboratories).

To localize CENP-E in RPE-1 cells, the cells were seeded onto 35 mm glass-bottom culture dishes (MatTek) for 24 h. Then cells were fixed with 4% PFA/PHEM buffer for 5 min at RT, then permeabilized with 0.1% Triton X-100/PHEM buffer for 2 min at RT and rinsed once with PHEM buffer. Cells were blocked for 30 min in antibody dilution solution (AbDil, 4% BSA, 0.1% Triton X-100, 0.1% sodium azide in PBS) and incubated with primary antibodies in AbDil overnight at 4°C. After washing for three times for 5 min each with PBST, cells were incubated with secondary antibodies in AbDil for 1 h at RT. The cells were washed three times for 5 min each with PBST, and DNA was stained with 100 ng/ml DAPI/PBS for 10 min at RT. After washing with PBS, cells were mounted with VECTASHIELD Mounting Medium (Vector Laboratories).

To localize CENP-E, CENP-C and CENP-T in RPE-1, K562, U2OS, A549, TIG-3, HeLa Kyoto, HT1080, OVCAR-3, HT29 or HCC1806 cells in Figure 6B-D, cells were treated with 50 μM monastrol (Selleckchem) for 2 h, then each cell line was collected and mixed with RPE-1 mScarlet-CENP-A cells as an internal control. Each mixed cell sample was cytospan onto coverslips (MATSUNAMI), and the cells were fixed and stained as described above. The CENP-E, CENP-C and CENP-T signal intensities at mitotic kinetochores in RPE-1, K562, U2OS, A549, TIG-3, HeLa Kyoto, HT1080, OVCAR-3, HT29 or HCC1806 cells were normalized with those signal intensities in mScarlet-CENP-A RPE-1 cells, which were in each sample.

Immunofluorescence images were captured by Nikon Eclipse Ti inverted microscope (Nikon) with a spinning disk confocal unit CSU-W1 (Yokogawa) controlled with NIS-elements (v5.42.01, Nikon) with an objective lens (Nikon; PlanApo VC 60×/1.40 or Lambda 100×/1.45 NA) and an Orca-Fusion BT (Hamamatsu Photonics) sCMOS camera. The images were acquired with Z-stacks at intervals of 0.2 or 0.3-µm. Maximum intensity projections (MIPs) of the Z-stack were generated using Fiji (v1.53)^86^ for display and analysis. All antibodies used in this study were listed in the Key resource table.

#### Lentivirus preparation

For large-scale lentiviral preparation, Lenti-X 293T cells (Takara Bio) were transfected with a transfection mix containing X-tremeGENE 9 DNA Transfection Reagent (Roche), pCMV-VSV-G (a gift from Robert Weinberg, Addgene plasmid, #8454),^89^ psPAX2 (a gift from Didier Trono, Addgene plasmid, #12260), and pooled sgRNA library plasmids^37,38^ in Opti-MEM I Reduced Serum Medium (Gibco). The culture medium was changed to a fresh medium at 16 h after the transfection. At 48 h after the transfection, the supernatant of the cell culture medium containing viruses was collected, filtered through a 0.45μm filter (Pall Corporation), and stored at −80°C until use.

#### Genome-Wide CRISPR Screening

The lentivirus pool containing the genome-wide sgRNA library was used to transduce 6×10^8^ K562 WT or CENP-C**^Δ^**^M12BD^ cells with 10 μg/ml polybrene (Merck) to maintain 1,000-fold coverage of the sgRNA library, as previously described ^90^. Three days after lentiviral transduction, the cells were passaged into fresh media supplemented with 3 μg/ml puromycin (Takara Bio) and selected for another 3 days. Collected 8×10^7^ cells were frozen at the endpoint of puromycin selection to assess initial sgRNA library representation (sgRNA^Day0^ abundance). The cells were allowed to grow with 1.5 μg/ml puromycin for 2 days then passaged with non-puromycin fresh medium every 2 days until 14 population doublings. Collected 8×10^7^ cells were frozen at the endpoint of the screen to assess the final sgRNA library representation (sgRNA^Day14^ abundance). The genomic DNA of Day 0 and Day 14 cells were extracted using the QIAamp DNA Blood Maxiprep Kit (Qiagen) according to the manufacturer’s instructions. For each extraction, the integrated sgRNAs were PCR-amplified for deep sequencing. Base calls were performed by the instrument control software and further processed using the Offline Base Caller (Illumina) v.1.9.4.

sgRNA sequences were counted and analyzed using MAGeCK v0.5.9.2,^91^ with a gene test false discovery rate (FDR) threshold of 0.05 and a list of control and intergenic sgRNAs for normalization and generation of the null distribution. The median of sgRNA scores was selected to represent the gene score. For comparison of gene knockout effect between K562 WT and CENP-C**^Δ^**^M12BD^ cells, genes with FDR < 0.1 were picked for further analysis (Table S1). A gene with a median fold change (log2) < -1 was identified as an essential gene, and a gene with a median fold change (log2) > -1 was identified as a nonessential gene. 375 candidate genes for non-essential in K562 WT cells but essential in K562 CENP-C**^Δ^**^M12BD^ cells were preranked based on the magnitude of differences of fold changes between K562 WT and CENP-C**^Δ^**^M12BD^ cells, in descending order. The preranked genes were analyzed with GSEA version 4.2.3 (Broad Institute) or R version 4.3.1 and all plots were generated in either base R, using the R corrplot package, or in Prism Version 10.1.2 (GraphPad).

#### RT-PCR

To isolate total RNA from K562 cells, 5×10^6^ cells were collected by centrifugation and suspended in TRIzol Reagent (Thermo Fisher Scientific), according to the manufacturer’s instructions. Using the isolated total RNA, CENP-C cDNA was reverse transcribed using SuperScript IV Reverse Transcriptase (Thermo Fisher Scientific), according to the manufacturer’s instructions. The cDNA was applied for PCR to examine the exon 2-4 deletion. The primes used are listed in Table S3.

#### Real-Time PCR

Total RNA was extracted from K562 cells with TRIzol (Thermo Fisher Scientific) as above. Relative mRNA expression levels were examined using iTaq Universal SYBR Green One-Step Kit (Bio-Rad) with QuantStudio 3 (Thermo Fisher Scientific). The Ct values of each gene were normalized with *GAPDH* Ct values in each sample. Data was collected and analyzed by the QuantStudio Design & Analysis Software v1.5.1 (Thermo Fisher Scientific). Primers used for Real-Time PCR were listed in Table S3.

#### Flow cytometry

5×10^6^ K562 cells were cultured with 20 μM 5-bromo-2-deoxyuridine (BrdU, Sigma-Aldrich) for 20 min. The cells were collected, washed with ice-cold PBS, fixed with ice-cold 70% ethanol, and stored at -20°C until use. The fixed cells were washed with 1% BSA/PBS and incubated with 4 M HCl/0.5% Triton X-100 for 30 min at RT. Following washing 3 times with 1% BSA/PBS, the cells were incubated with an anti-BrdU antibody (BD Biosciences) for 1 h at RT. Following washing twice with 1% BSA/PBS, the cells were incubated with 1:20 anti-mouse IgG-FITC (Jackson ImmunoResearch) in 1% BSA/PBS for 30 min at RT. After a wash with 1% BSA/PBS, the cells were incubated with 10 μg/ml propidium iodide (Sigma-Aldrich) in 1% BSA/PBS overnight at 4°C. The stained cells were subjected to flow cytometry analysis using BD FACS Canto II Flow Cytometer (BD Biosciences), and BD FACSDiva 9.0 Software (BD Biosciences).

#### Cell viability assay

Cell viability was assessed using RealTime-Glo MT Cell Viability Assay (Promega). K562 cells were transferred to 96-well Flat Clear Bottom White Polystyrene TC-treated Microplates (Corning) at the indicated time points. The cells were treated with RealTime-Glo reagents (Promega) and incubated for 1 h at 37°C. Luminescence was measured using GloMax Discover Microplate Reader (Promega) and GloMax Navigator (Promega). RPE-1, U2OS, A549, TIG-3, HeLa Kyoto, HT1080, OVCAR-3, HT29 or HCC1806 cells were seeded into 96-well Flat Clear Bottom White Polystyrene TC-treated Microplates (Corning) at the concentration of 1250 cells/well. Following culture for 3 days, the cells were treated with RealTime-Glo reagents (Promega) and incubated for 1 h at 37°C. Luminescence was measured using GloMax Discover Microplate Reader (Promega) and GloMax Navigator (Promega).

#### siRNA treatment

3 μl Lipofectamine RNAiMAX Reagent (Thermo Fisher Scientific) and 10 pmol siRNA were diluted in 125 μl Opti-MEM I Reduced Serum Medium (Gibco) separately. Diluted siRNA and diluted Lipofectamine RNAiMAX Reagent were mixed at 1:1 ratio and were incubated for 5 min at RT. The 250 μl siRNA-lipid complex was added into a 6-well plate (IWAKI) or 35 mm glass-bottom culture dishes (MatTek), then 5 – 7.5×10^4^ cells were added into the plate or 35 mm dish. Cells were incubated for 1-3 days at 37°C before analyzing. For the cell viability assay, 20 μl siRNA-lipid complex was added into 96-well Flat Clear Bottom White Polystyrene TC-treated Microplates (Corning), then 1250 cells/80 μl were added into the plate. Cells were incubated for 3 days at 37°C before analyzing. The sequences of siRNA used in this study are listed in Table S3.

#### Live-cell imaging

For live-cell imaging, we used RPE-1 cells expressing mScarlet-CENP-A, GFP-H2A, and Flag-fused CENP-C^WT^ or CENP-C^ΔM12BD^ (RPE-1 mScarlet-CENP-A GFP-H2A Flag-CENP-C^WT^, RPE-1 mScarlet-CENP-A GFP-H2A Flag-CENP-C^ΔM12BD^).^23^ 1×10^5^ cells were treated with siRNA as described in the above section and plated onto 35 mm glass-bottom culture dishes (MatTek). After incubation at 37°C for 24 h, cells were switched to phenol red-free culture medium (phenol red-free DMEM (Nakalai Tesque), supplemented with 20% fetal bovine serum (Sigma-Aldrich), 25 mM HEPES (HEPES Buffer Solution, Nacalai Tesque), 2 mM L-glutamine (Sigma-Aldrich) and treated with SiR-Tubulin (1:5,000) (CYTOSKELETON) for 4 h. Then, samples were sealed with mineral oil (Sigma-Aldrich) before the start of imaging. To quantify mitotic timing in Figure 3G, the images were captured every 2 min by Olympus IX71 microscope (Olympus) with DeltaVision Elite imaging system (GE healthcare) equipped with a PlanApo N OSC 60×/1.40 NA oil immersion objective lens (Olympus) and a CoolSNAP HQ2 CCD camera (Photometrics) controlled with built-in SoftWoRx software (version 5.5) in a temperature-controlled room at 37°C. A Z-series of 7 sections with 2 µm increments was acquired. The Z-series was then projected using MIP for analysis by SoftWoRx. To quantify peripheral chromosome congression in Figures 5C and 5F, the images were captured every 1 min by Nikon Eclipse Ti inverted microscope (Nikon) with a spinning disk confocal unit CSU-W1 (Yokogawa) controlled with NIS-elements (v5.42.01, Nikon) with an objective lens (Nikon; PlanApo VC 60×/1.40) and an Orca-Fusion BT (Hamamatsu Photonics) sCMOS camera. The images were acquired with Z-stacks at intervals of 2 µm. MIP of the Z-stack were generated using NIS-elements Imaging Software (v5.42.01, Nikon). Acquired time-lapse images were processed by using Fiji software (v1.53).^86^

### QUANTIFICATION AND STATISTICAL ANALYSIS

The fluorescence signal intensities of CENP-C, CENP-T, DSN1, KNL1, HEC1, CENP-E, ZW10 and mScarlet-DSN1 on kinetochores were quantified using Imaris v9.0.2. (Bitplane). Data processing was carried out using Microsoft Excel and GraphPad Prism 10.1.2 (GraphPad), and p-values were calculated using two-tailed Student’s t-test. Cell viability was measured by luminescence using GloMax Discover Microplate Reader (Promega) and GloMax Navigator (Promega). Data processing was carried out using Microsoft Excel and GraphPad Prism 10.1.2 (GraphPad), and p-values were calculated using two-tailed Student’s t-test. Each experiment was repeated at least three times when not mentioned in figure legends, and representative data of replicates were presented.

## Supplementary Information

The article contains 7 Supplementary Figures, 3 Supplementary Tables and 4 Supplementary movies.

## Key resources table

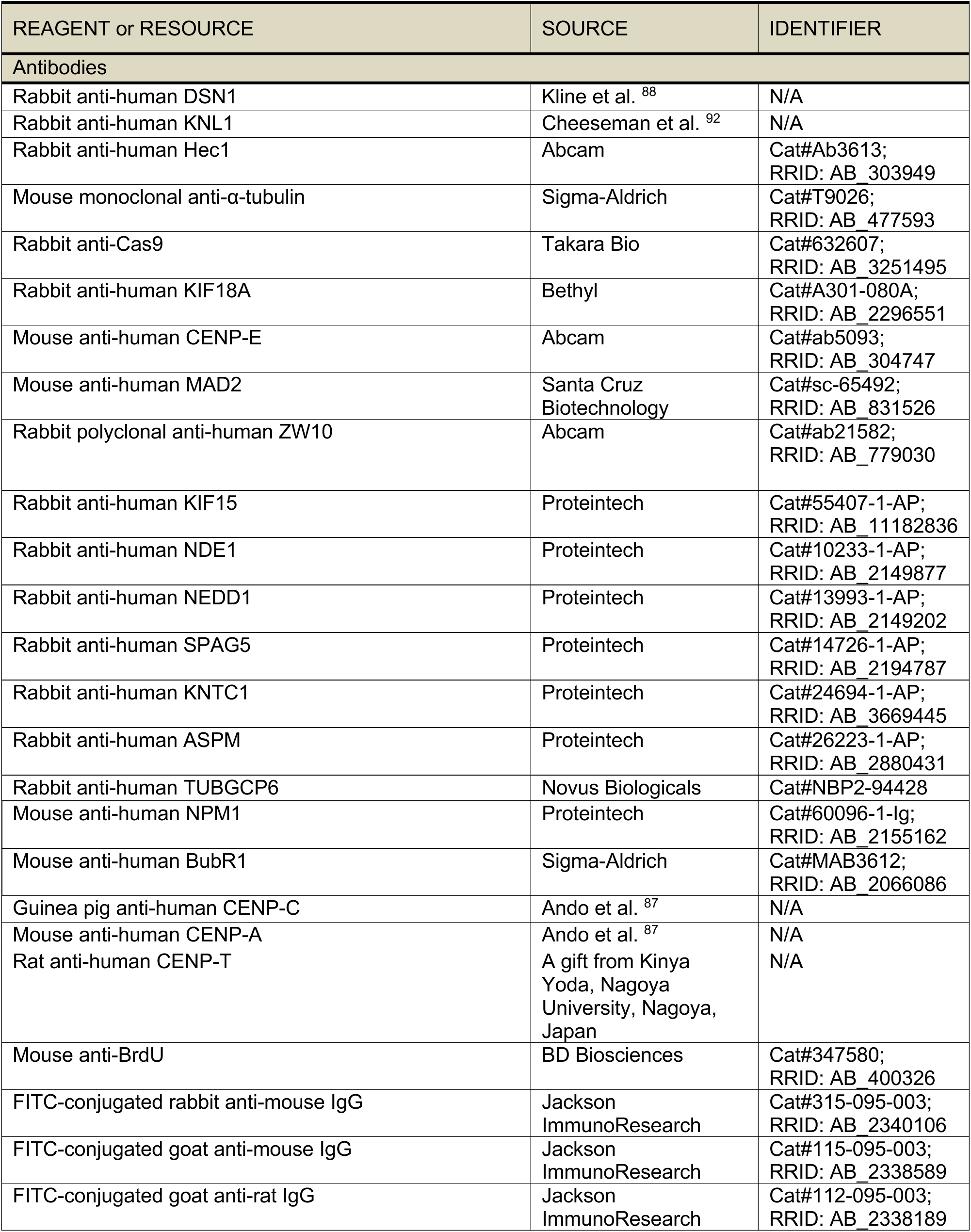
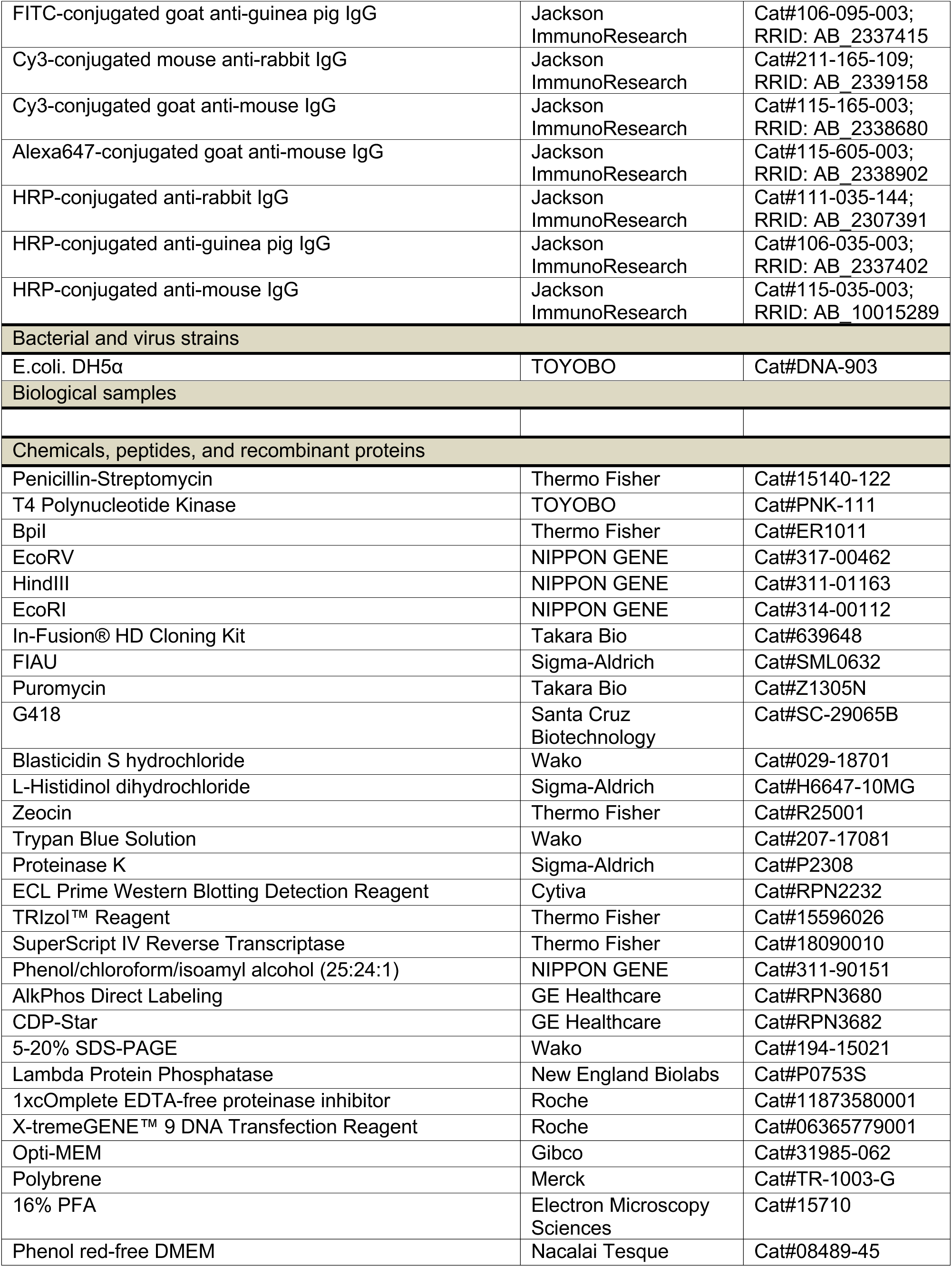
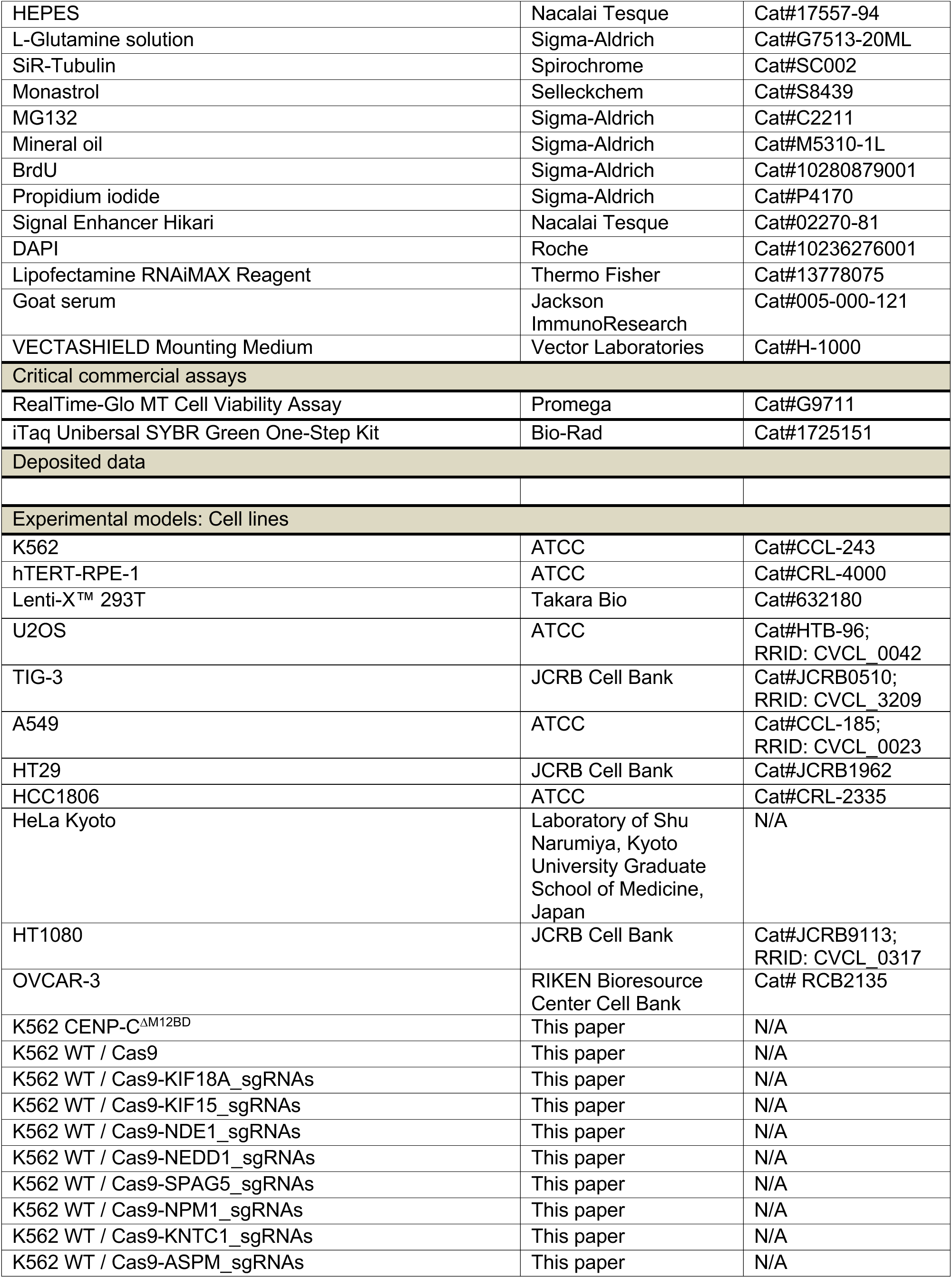
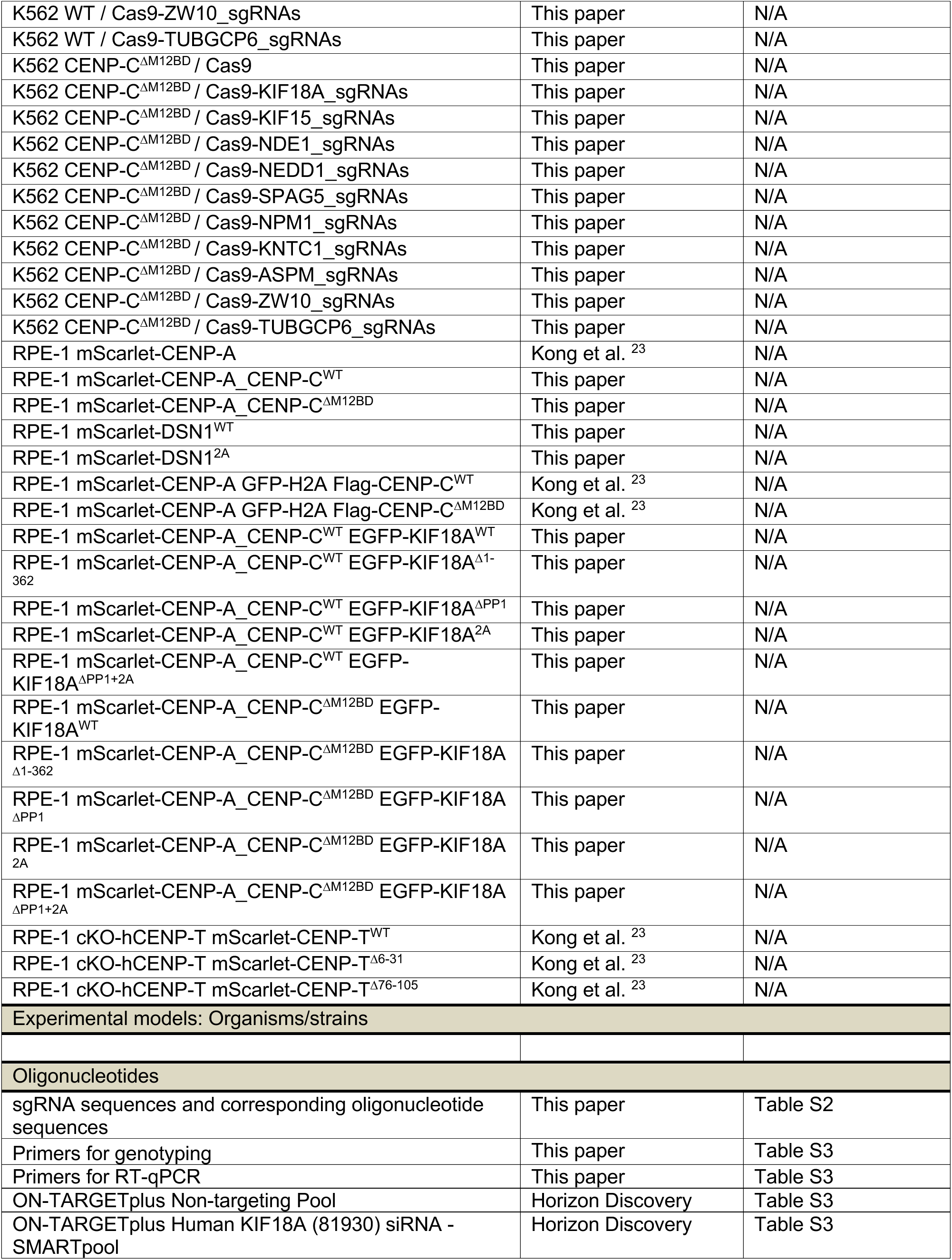
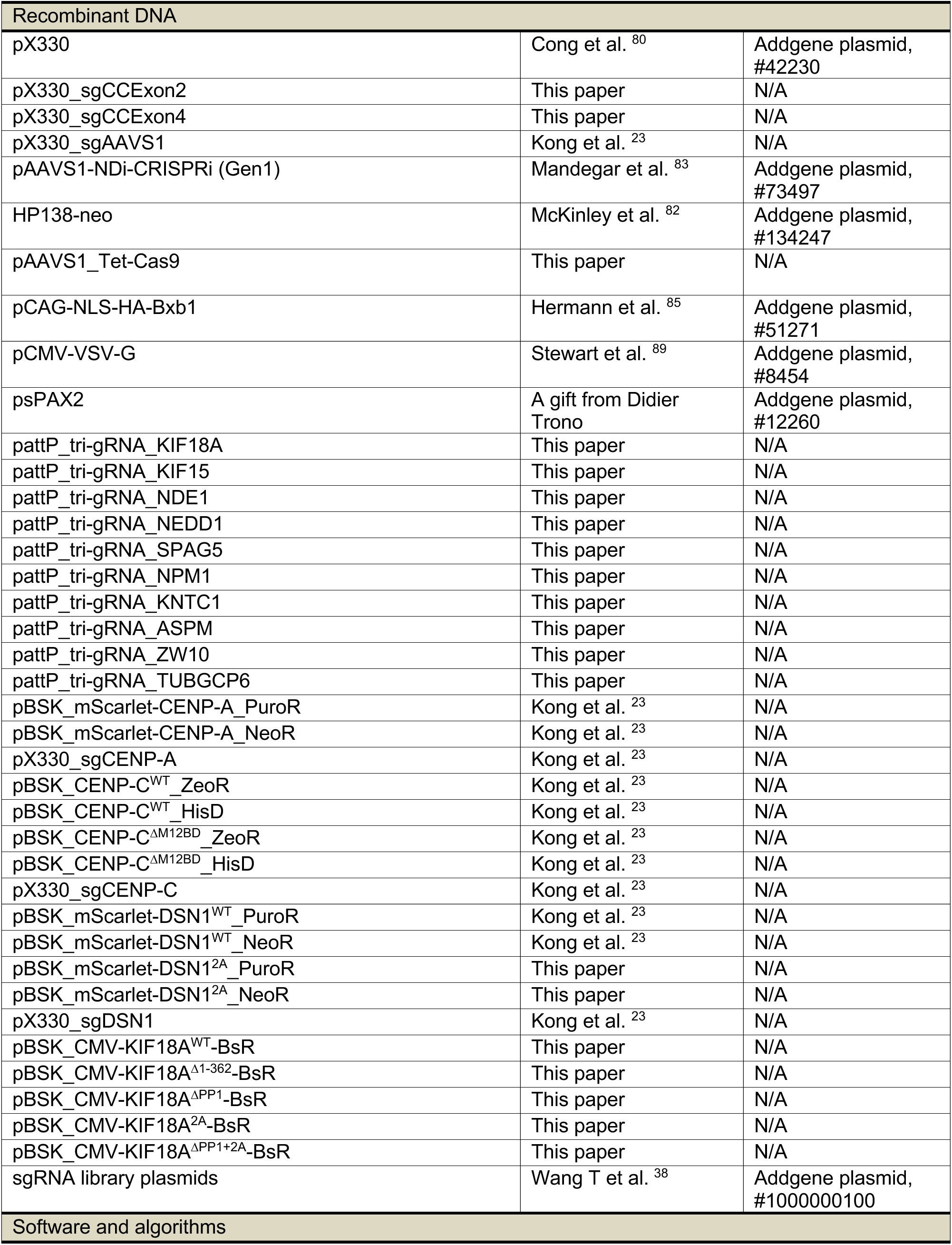
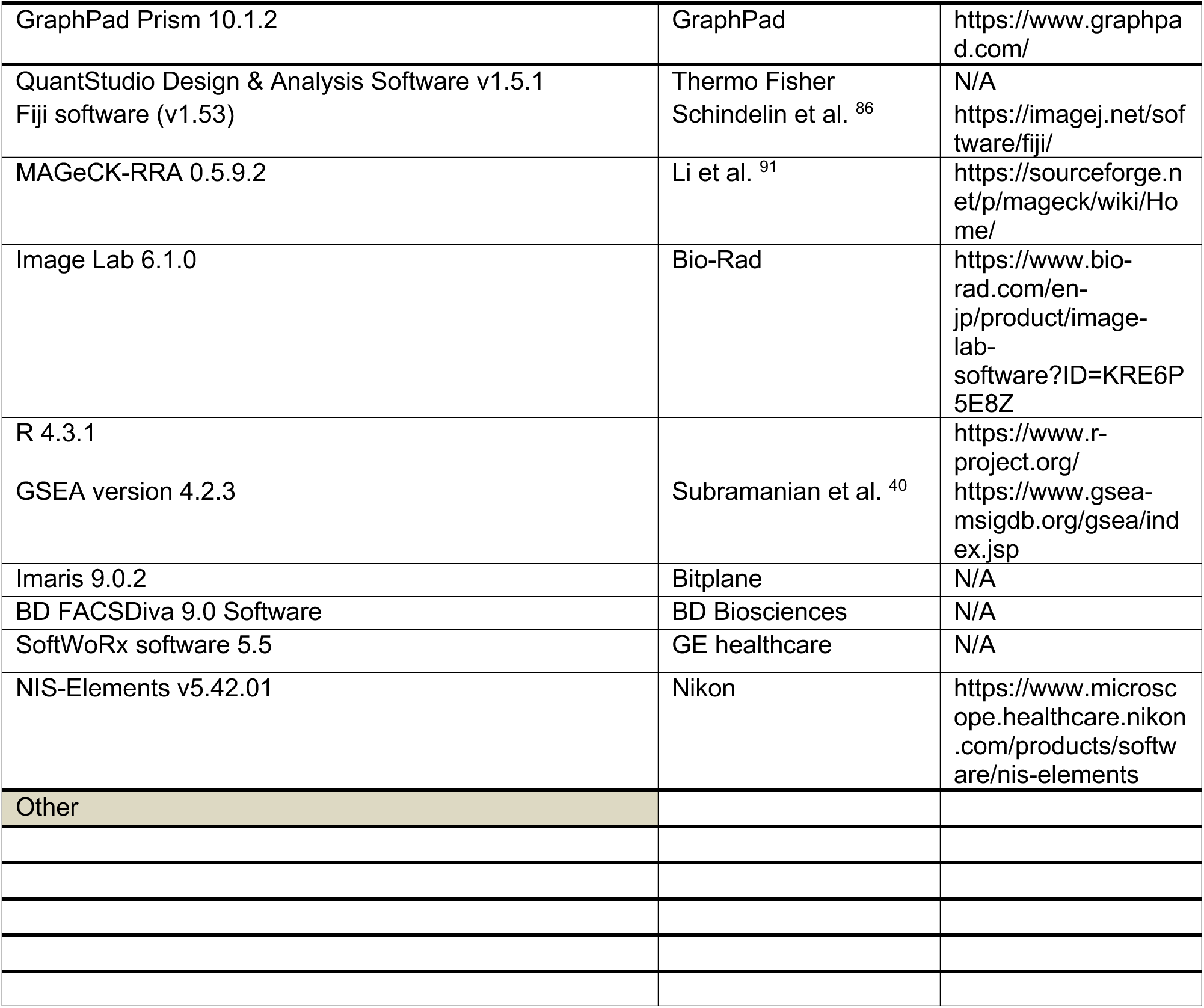

| REAGENT or RESOURCE | SOURCE | IDENTIFIER |
| --- | --- | --- |
| **Antibodies** |  |  |
| Rabbit anti-human DSN1 | Kline et al. <sup>88</sup> | N/A |
| Rabbit anti-human KNL1 | Cheeseman et al. <sup>92</sup> | N/A |
| Rabbit anti-human Hec1 | Abcam | Cat#Ab3613;<br>RRID: AB_303949 |
| Mouse monoclonal anti-α-tubulin | Sigma-Aldrich | Cat#T9026;<br>RRID: AB_477593 |
| Rabbit anti-Cas9 | Takara Bio | Cat#632607;<br>RRID: AB_3251495 |
| Rabbit anti-human KIF18A | Bethyl | Cat#A301-080A;<br>RRID: AB_2296551 |
| Mouse anti-human CENP-E | Abcam | Cat#ab5093;<br>RRID: AB_304747 |
| Mouse anti-human MAD2 | Santa Cruz<br>Biotechnology | Cat#sc-65492;<br>RRID: AB_831526 |
| Rabbit polyclonal anti-human ZW10 | Abcam | Cat#ab21582;<br>RRID: AB_779030 |
| Rabbit anti-human KIF15 | Proteintech | Cat#55407-1-AP;<br>RRID: AB_11182836 |
| Rabbit anti-human NDE1 | Proteintech | Cat#10233-1-AP;<br>RRID: AB_2149877 |
| Rabbit anti-human NEDD1 | Proteintech | Cat#13993-1-AP;<br>RRID: AB_2149202 |
| Rabbit anti-human SPAG5 | Proteintech | Cat#14726-1-AP;<br>RRID: AB_2194787 |
| Rabbit anti-human KNTC1 | Proteintech | Cat#24694-1-AP;<br>RRID: AB_3669445 |
| Rabbit anti-human ASPM | Proteintech | Cat#26223-1-AP;<br>RRID: AB_2880431 |
| Rabbit anti-human TUBGCP6 | Novus Biologicals | Cat#NBP2-94428 |
| Mouse anti-human NPM1 | Proteintech | Cat#60096-1-Ig;<br>RRID: AB_2155162 |
| Mouse anti-human BubR1 | Sigma-Aldrich | Cat#MAB3612;<br>RRID: AB_2066086 |
| Guinea pig anti-human CENP-C | Ando et al. <sup>87</sup> | N/A |
| Mouse anti-human CENP-A | Ando et al. <sup>87</sup> | N/A |
| Rat anti-human CENP-T | A gift from Kinuya<br>Yoda, Nagoya<br>University, Nagoya,<br>Japan | N/A |
| Mouse anti-BrdU | BD Biosciences | Cat#347580;<br>RRID: AB_400326 |
| FITC-conjugated rabbit anti-mouse IgG | Jackson<br>ImmunoResearch | Cat#315-095-003;<br>RRID: AB_2340106 |
| FITC-conjugated goat anti-mouse IgG | Jackson<br>ImmunoResearch | Cat#115-095-003;<br>RRID: AB_2338589 |
| FITC-conjugated goat anti-rat IgG | Jackson<br>ImmunoResearch | Cat#112-095-003;<br>RRID: AB_2338189 |
| FITC-conjugated goat anti-guinea pig IgG | Jackson ImmunoResearch | Cat#106-095-003; RRID: AB_2337415 |
| Cy3-conjugated mouse anti-rabbit IgG | Jackson ImmunoResearch | Cat#211-165-109; RRID: AB_2339158 |
| Cy3-conjugated goat anti-mouse IgG | Jackson ImmunoResearch | Cat#115-165-003; RRID: AB_2338680 |
| Alexa647-conjugated goat anti-mouse IgG | Jackson ImmunoResearch | Cat#115-605-003; RRID: AB_2338902 |
| HRP-conjugated anti-rabbit IgG | Jackson ImmunoResearch | Cat#111-035-144; RRID: AB_2307391 |
| HRP-conjugated anti-guinea pig IgG | Jackson ImmunoResearch | Cat#106-035-003; RRID: AB_2337402 |
| HRP-conjugated anti-mouse IgG | Jackson ImmunoResearch | Cat#115-035-003; RRID: AB_10015289 |
| Bacterial and virus strains |  |  |
| E.coli. DH5α | TOYOBO | Cat#DNA-903 |
| Biological samples |  |  |
| Chemicals, peptides, and recombinant proteins |  |  |
| Penicillin-Streptomycin | Thermo Fisher | Cat#15140-122 |
| T4 Polynucleotide Kinase | TOYOBO | Cat#PNK-111 |
| Bpil | Thermo Fisher | Cat#ER1011 |
| EcoRV | NIPPON GENE | Cat#317-00462 |
| HindIII | NIPPON GENE | Cat#311-01163 |
| EcoRI | NIPPON GENE | Cat#314-00112 |
| In-Fusion® HD Cloning Kit | Takara Bio | Cat#639648 |
| FIAU | Sigma-Aldrich | Cat#SML0632 |
| Puromycin | Takara Bio | Cat#Z1305N |
| G418 | Santa Cruz Biotechnology | Cat#SC-29065B |
| Blasticidin S hydrochloride | Wako | Cat#029-18701 |
| L-Histidinol dihydrochloride | Sigma-Aldrich | Cat#H6647-10MG |
| Zeocin | Thermo Fisher | Cat#R25001 |
| Trypan Blue Solution | Wako | Cat#207-17081 |
| Proteinase K | Sigma-Aldrich | Cat#P2308 |
| ECL Prime Western Blotting Detection Reagent | Cytiva | Cat#RPN2232 |
| TRIzol™ Reagent | Thermo Fisher | Cat#15596026 |
| SuperScript IV Reverse Transcriptase | Thermo Fisher | Cat#18090010 |
| Phenol/chloroform/isoamyl alcohol (25:24:1) | NIPPON GENE | Cat#311-90151 |
| AlkPhos Direct Labeling | GE Healthcare | Cat#RPN3680 |
| CDP-Star | GE Healthcare | Cat#RPN3682 |
| 5-20% SDS-PAGE | Wako | Cat#194-15021 |
| Lambda Protein Phosphatase | New England Biolabs | Cat#P0753S |
| 1xComplete EDTA-free proteinase inhibitor | Roche | Cat#11873580001 |
| X-tremeGENE™ 9 DNA Transfection Reagent | Roche | Cat#06365779001 |
| Opti-MEM | Gibco | Cat#31985-062 |
| Polybrene | Merck | Cat#TR-1003-G |
| 16% PFA | Electron Microscopy Sciences | Cat#15710 |
| Phenol red-free DMEM | Nacalai Tesque | Cat#08489-45 |
| HEPES | Nacalai Tesque | Cat#17557-94 |
| L-Glutamine solution | Sigma-Aldrich | Cat#G7513-20ML |
| SiR-Tubulin | Spirochrome | Cat#SC002 |
| Monastrol | Selleckchem | Cat#S8439 |
| MG132 | Sigma-Aldrich | Cat#C2211 |
| Mineral oil | Sigma-Aldrich | Cat#M5310-1L |
| BrdU | Sigma-Aldrich | Cat#10280879001 |
| Propidium iodide | Sigma-Aldrich | Cat#P4170 |
| Signal Enhancer Hikari | Nacalai Tesque | Cat#02270-81 |
| DAPI | Roche | Cat#10236276001 |
| Lipofectamine RNAiMAX Reagent | Thermo Fisher | Cat#13778075 |
| Goat serum | Jackson ImmunoResearch | Cat#005-000-121 |
| VECTASHIELD Mounting Medium | Vector Laboratories | Cat#H-1000 |
| Critical commercial assays |  |  |
| RealTime-Glo MT Cell Viability Assay | Promega | Cat#G9711 |
| iTaq Universal SYBR Green One-Step Kit | Bio-Rad | Cat#1725151 |
| Deposited data |  |  |
| Experimental models: Cell lines |  |  |
| K562 | ATCC | Cat#CCL-243 |
| hTERT-RPE-1 | ATCC | Cat#CRL-4000 |
| Lenti-X™ 293T | Takara Bio | Cat#632180 |
| U2OS | ATCC | Cat#HTB-96;<br>RRID: CVCL_0042 |
| TIG-3 | JCRB Cell Bank | Cat#JCRB0510;<br>RRID: CVCL_3209 |
| A549 | ATCC | Cat#CCL-185;<br>RRID: CVCL_0023 |
| HT29 | JCRB Cell Bank | Cat#JCRB1962 |
| HCC1806 | ATCC | Cat#CRL-2335 |
| HeLa Kyoto | Laboratory of Shu Narumiya, Kyoto University Graduate School of Medicine, Japan | N/A |
| HT1080 | JCRB Cell Bank | Cat#JCRB9113;<br>RRID: CVCL_0317 |
| OVCAR-3 | RIKEN Bioresource Center Cell Bank | Cat# RCB2135 |
| K562 CENP-C<sup>ΔM12BD</sup> | This paper | N/A |
| K562 WT / Cas9 | This paper | N/A |
| K562 WT / Cas9-KIF18A_sgRNAs | This paper | N/A |
| K562 WT / Cas9-KIF15_sgRNAs | This paper | N/A |
| K562 WT / Cas9-NDE1_sgRNAs | This paper | N/A |
| K562 WT / Cas9-NEDD1_sgRNAs | This paper | N/A |
| K562 WT / Cas9-SPAG5_sgRNAs | This paper | N/A |
| K562 WT / Cas9-NPM1_sgRNAs | This paper | N/A |
| K562 WT / Cas9-KNTC1_sgRNAs | This paper | N/A |
| K562 WT / Cas9-ASPM_sgRNAs | This paper | N/A |
| K562 WT / Cas9-ZW10_sgRNAs | This paper | N/A |
| K562 WT / Cas9-TUBGCP6_sgRNAs | This paper | N/A |
| K562 CENP-C<sup>ΔM12BD</sup> / Cas9 | This paper | N/A |
| K562 CENP-C<sup>ΔM12BD</sup> / Cas9-KIF18A_sgRNAs | This paper | N/A |
| K562 CENP-C<sup>ΔM12BD</sup> / Cas9-KIF15_sgRNAs | This paper | N/A |
| K562 CENP-C<sup>ΔM12BD</sup> / Cas9-NDE1_sgRNAs | This paper | N/A |
| K562 CENP-C<sup>ΔM12BD</sup> / Cas9-NEDD1_sgRNAs | This paper | N/A |
| K562 CENP-C<sup>ΔM12BD</sup> / Cas9-SPAG5_sgRNAs | This paper | N/A |
| K562 CENP-C<sup>ΔM12BD</sup> / Cas9-NPM1_sgRNAs | This paper | N/A |
| K562 CENP-C<sup>ΔM12BD</sup> / Cas9-KNTC1_sgRNAs | This paper | N/A |
| K562 CENP-C<sup>ΔM12BD</sup> / Cas9-ASPM_sgRNAs | This paper | N/A |
| K562 CENP-C<sup>ΔM12BD</sup> / Cas9-ZW10_sgRNAs | This paper | N/A |
| K562 CENP-C<sup>ΔM12BD</sup> / Cas9-TUBGCP6_sgRNAs | This paper | N/A |
| RPE-1 mScarlet-CENP-A | Kong et al. <sup>23</sup> | N/A |
| RPE-1 mScarlet-CENP-A_CENP-C<sup>WT</sup> | This paper | N/A |
| RPE-1 mScarlet-CENP-A_CENP-C<sup>ΔM12BD</sup> | This paper | N/A |
| RPE-1 mScarlet-DSN1<sup>WT</sup> | This paper | N/A |
| RPE-1 mScarlet-DSN1<sup>2A</sup> | This paper | N/A |
| RPE-1 mScarlet-CENP-A GFP-H2A Flag-CENP-C<sup>WT</sup> | Kong et al. <sup>23</sup> | N/A |
| RPE-1 mScarlet-CENP-A GFP-H2A Flag-CENP-C<sup>ΔM12BD</sup> | Kong et al. <sup>23</sup> | N/A |
| RPE-1 mScarlet-CENP-A_CENP-C<sup>WT</sup> EGFP-KIF18A<sup>WT</sup> | This paper | N/A |
| RPE-1 mScarlet-CENP-A_CENP-C<sup>WT</sup> EGFP-KIF18A<sup>Δ1-362</sup> | This paper | N/A |
| RPE-1 mScarlet-CENP-A_CENP-C<sup>WT</sup> EGFP-KIF18A<sup>ΔPP1</sup> | This paper | N/A |
| RPE-1 mScarlet-CENP-A_CENP-C<sup>WT</sup> EGFP-KIF18A<sup>2A</sup> | This paper | N/A |
| RPE-1 mScarlet-CENP-A_CENP-C<sup>WT</sup> EGFP-KIF18A<sup>ΔPP1+2A</sup> | This paper | N/A |
| RPE-1 mScarlet-CENP-A_CENP-C<sup>ΔM12BD</sup> EGFP-KIF18A<sup>WT</sup> | This paper | N/A |
| RPE-1 mScarlet-CENP-A_CENP-C<sup>ΔM12BD</sup> EGFP-KIF18A<sup>Δ1-362</sup> | This paper | N/A |
| RPE-1 mScarlet-CENP-A_CENP-C<sup>ΔM12BD</sup> EGFP-KIF18A<sup>ΔPP1</sup> | This paper | N/A |
| RPE-1 mScarlet-CENP-A_CENP-C<sup>ΔM12BD</sup> EGFP-KIF18A<sup>2A</sup> | This paper | N/A |
| RPE-1 mScarlet-CENP-A_CENP-C<sup>ΔM12BD</sup> EGFP-KIF18A<sup>ΔPP1+2A</sup> | This paper | N/A |
| RPE-1 cKO-hCENP-T mScarlet-CENP-T<sup>WT</sup> | Kong et al. <sup>23</sup> | N/A |
| RPE-1 cKO-hCENP-T mScarlet-CENP-T<sup>Δ6-31</sup> | Kong et al. <sup>23</sup> | N/A |
| RPE-1 cKO-hCENP-T mScarlet-CENP-T<sup>Δ76-105</sup> | Kong et al. <sup>23</sup> | N/A |
| Experimental models: Organisms/strains |  |  |
| Oligonucleotides |  |  |
| sgRNA sequences and corresponding oligonucleotide sequences | This paper | Table S2 |
| Primers for genotyping | This paper | Table S3 |
| Primers for RT-qPCR | This paper | Table S3 |
| ON-TARGETplus Non-targeting Pool | Horizon Discovery | Table S3 |
| ON-TARGETplus Human KIF18A (81930) siRNA - SMARTpool | Horizon Discovery | Table S3 |

| Recombinant DNA |  |  |
| --- | --- | --- |
| pX330 | Cong et al. <sup>80</sup> | Addgene plasmid, #42230 |
| pX330_sgCCExon2 | This paper | N/A |
| pX330_sgCCExon4 | This paper | N/A |
| pX330_sgAAVS1 | Kong et al. <sup>23</sup> | N/A |
| pAAVS1-NDi-CRISPRi (Gen1) | Mandegar et al. <sup>83</sup> | Addgene plasmid, #73497 |
| HP138-neo | McKinley et al. <sup>82</sup> | Addgene plasmid, #134247 |
| pAAVS1_Tet-Cas9 | This paper | N/A |
| pCAG-NLS-HA-Bxb1 | Hermann et al. <sup>85</sup> | Addgene plasmid, #51271 |
| pCMV-VSV-G | Stewart et al. <sup>89</sup> | Addgene plasmid, #8454 |
| psPAX2 | A gift from Didier Trono | Addgene plasmid, #12260 |
| pattP_tri-gRNA_KIF18A | This paper | N/A |
| pattP_tri-gRNA_KIF15 | This paper | N/A |
| pattP_tri-gRNA_NDE1 | This paper | N/A |
| pattP_tri-gRNA_NEDD1 | This paper | N/A |
| pattP_tri-gRNA_SPAG5 | This paper | N/A |
| pattP_tri-gRNA_NPM1 | This paper | N/A |
| pattP_tri-gRNA_KNTC1 | This paper | N/A |
| pattP_tri-gRNA_ASPM | This paper | N/A |
| pattP_tri-gRNA_ZW10 | This paper | N/A |
| pattP_tri-gRNA_TUBGCP6 | This paper | N/A |
| pBSK_mScarlet-CENP-A_PuroR | Kong et al. <sup>23</sup> | N/A |
| pBSK_mScarlet-CENP-A_NeoR | Kong et al. <sup>23</sup> | N/A |
| pX330_sgCENP-A | Kong et al. <sup>23</sup> | N/A |
| pBSK_CENP-C<sup>WT</sup>_ZeoR | Kong et al. <sup>23</sup> | N/A |
| pBSK_CENP-C<sup>WT</sup>_HisD | Kong et al. <sup>23</sup> | N/A |
| pBSK_CENP-C<sup>ΔM12BD</sup>_ZeoR | Kong et al. <sup>23</sup> | N/A |
| pBSK_CENP-C<sup>ΔM12BD</sup>_HisD | Kong et al. <sup>23</sup> | N/A |
| pX330_sgCENP-C | Kong et al. <sup>23</sup> | N/A |
| pBSK_mScarlet-DSN1<sup>WT</sup>_PuroR | Kong et al. <sup>23</sup> | N/A |
| pBSK_mScarlet-DSN1<sup>WT</sup>_NeoR | Kong et al. <sup>23</sup> | N/A |
| pBSK_mScarlet-DSN1<sup>2A</sup>_PuroR | This paper | N/A |
| pBSK_mScarlet-DSN1<sup>2A</sup>_NeoR | This paper | N/A |
| pX330_sgDSN1 | Kong et al. <sup>23</sup> | N/A |
| pBSK_CMV-KIF18A<sup>WT</sup>-BsR | This paper | N/A |
| pBSK_CMV-KIF18A<sup>Δ1-362</sup>-BsR | This paper | N/A |
| pBSK_CMV-KIF18A<sup>ΔPP1</sup>-BsR | This paper | N/A |
| pBSK_CMV-KIF18A<sup>2A</sup>-BsR | This paper | N/A |
| pBSK_CMV-KIF18A<sup>ΔPP1+2A</sup>-BsR | This paper | N/A |
| sgRNA library plasmids | Wang T et al. <sup>38</sup> | Addgene plasmid, #1000000100 |
| Software and algorithms |  |  |
| GraphPad Prism 10.1.2 | GraphPad | https://www.graphpad.com/ |
| QuantStudio Design & Analysis Software v1.5.1 | Thermo Fisher | N/A |
| Fiji software (v1.53) | Schindelin et al. <sup>86</sup> | https://imagej.net/software/fiji/ |
| MAGeCK-RRA 0.5.9.2 | Li et al. <sup>91</sup> | https://sourceforge.net/p/mageck/wiki/Home/ |
| Image Lab 6.1.0 | Bio-Rad | https://www.bio-rad.com/en-jp/product/image-lab-software?ID=KRE6P5E8Z |
| R 4.3.1 |  | https://www.r-project.org/ |
| GSEA version 4.2.3 | Subramanian et al. <sup>40</sup> | https://www.gsea-msigdb.org/gsea/index.jsp |
| Imaris 9.0.2 | Bitplane | N/A |
| BD FACSDiva 9.0 Software | BD Biosciences | N/A |
| SoftWoRx software 5.5 | GE healthcare | N/A |
| NIS-Elements v5.42.01 | Nikon | https://www.microscope.healthcare.nikon.com/products/software/nis-elements |
| Other |  |  |

## Legends for Supplementary Figures, Tables and Movies

**Figure S1.**
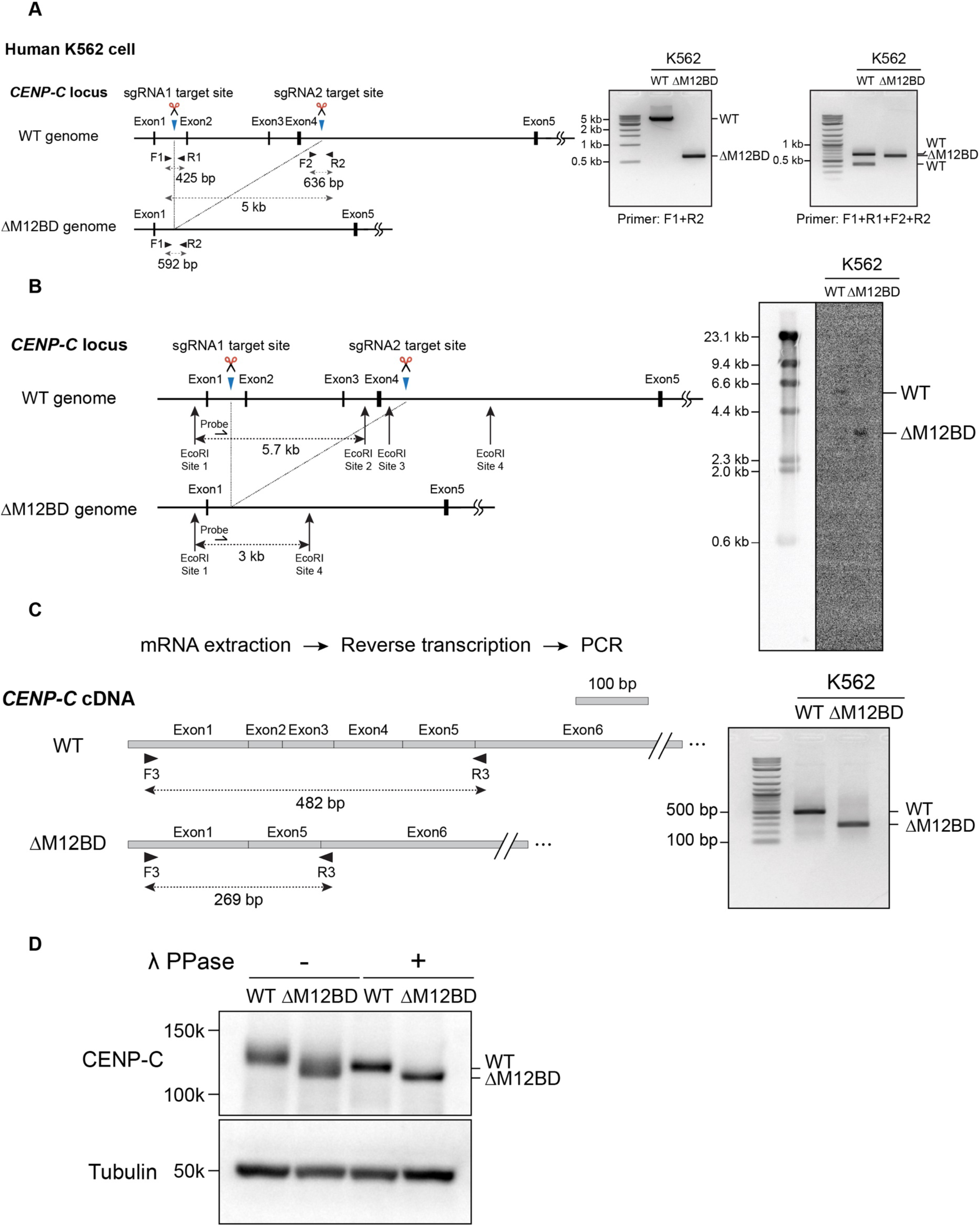
Generation of K562 CENP-C^ΔM12BD^ cells. (A) Schematic representation of the exon 2-4 deletion from human *CENP-C* gene locus in K562 cells. Using the CRISPR-Cas9 system with two sgRNAs, the exon 2-4 is deleted to generate K562 CENP-C^ΔM12BD^ cells. The position of sgRNAs and PCR primers for genotyping PCR are shown. The amplicon sizes are 425 bp (F1+R1) in K562 WT cells, 636 bp (F2+R2) in K562 WT cells, 592 bp (F1+R2) in K562 CENP-C^ΔM12BD^ cells. Results of genotyping PCR with K562 WT and CENP-C^ΔM12BD^ cells are shown. (B) Design for Southern blot analysis to detect a deletion of the exon 2-4 at the *CENP-C* gene locus in K562 CENP-C^ΔM12BD^ cells. EcoRI sites and the probe are shown. Purified genomic DNA from K562 WT or CENP-C^ΔM12BD^ cells was digested with EcoRI and run on an agarose gel. The DNA fragments on the gel were transferred to the nylon membrane and hybridized with the indicated probe. (C) RT-PCR to examine the expression and exon 2-4 deletion of mRNA for wild-type (WT) CENP-C and CENP-C^ΔM12BD^ in K562 cells. (D) Immunoblot analysis to detect WT CENP-C and CENP-C^ΔM12BD^ in each cell line. α-tubulin was probed as a loading control. As CENP-C is highly phosphorylated, CENP-C band was smeared. To detect the CENP-C protein band, PPase was used to dephosphorylate proteins.

**Figure S2.**
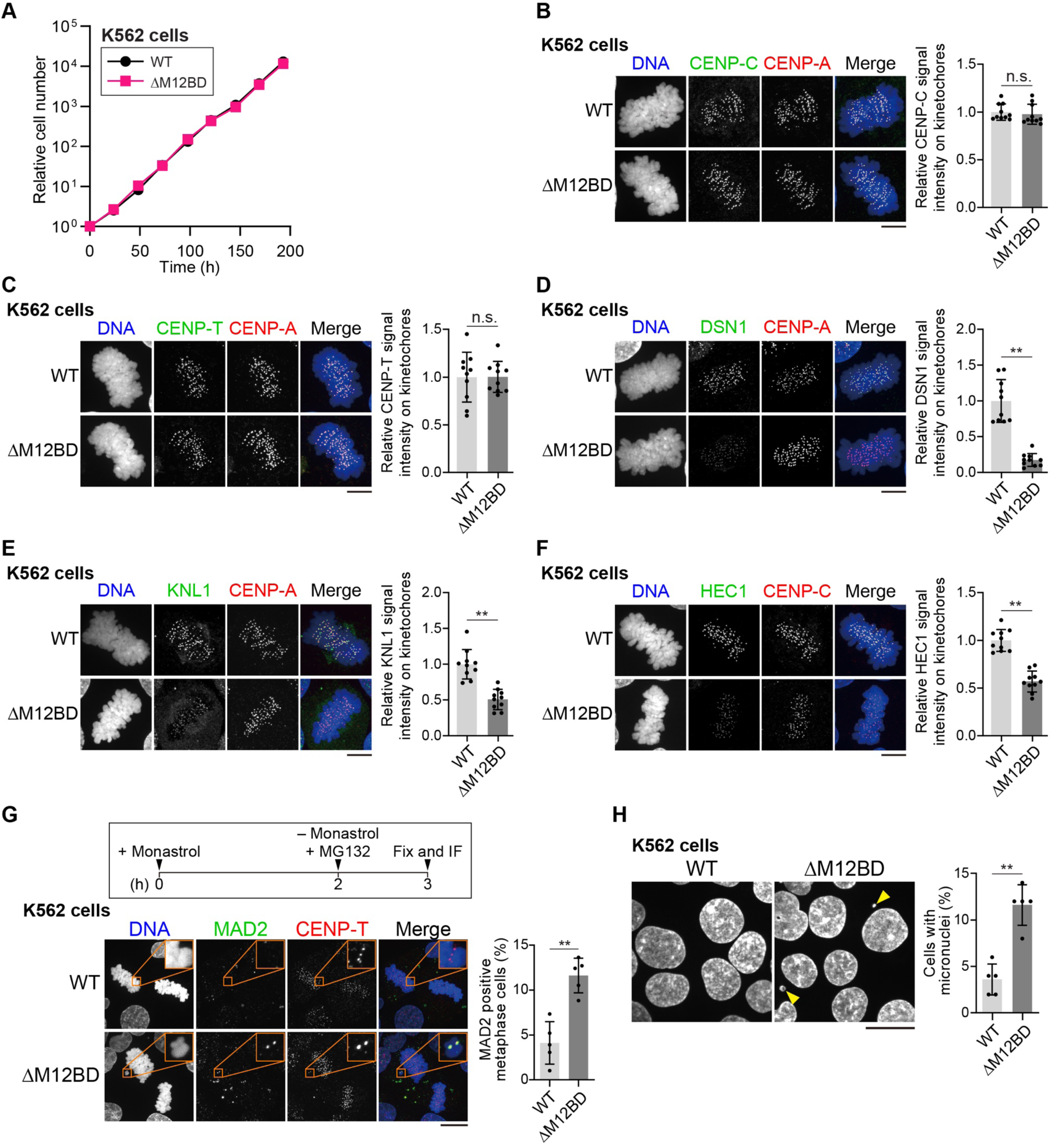
Characterization of K562 CENP-C^ΔM12BD^ cells. (A) The growth curve of K562 WT or CENP-C^ΔM12BD^ cells. The cell numbers were normalized to those at time 0 of each cell line. (B-F) CENP-C, CENP-T, DSN1, KNL1, and HEC1 localization in K562 WT or CENP-C^ΔM12BD^ cells. Each protein was stained with an antibody against the target protein (green). CENP-A was stained as a kinetochore marker (red) except for HEC1 staining. For HEC1 staining, CENP-C was stained as a kinetochore marker (red). DNA was stained with DAPI (blue). Scale bar, 10 μm. Signal intensities of target protein on mitotic kinetochores were quantified (Mean and SD, two-tailed Student’s t-test, K562 WT cells: n = 10 cells, K562 CENP-C^ΔM12BD^ cells: n = 10 cells; n.s., non-significant; \*\**p* < 0.01). (G) MAD2 staining (green) in K562 WT or CENP-C^ΔM12BD^ cells. Cells were treated with Monastrol for 2 h and after washout of Monastrol, cells were treated with MG132 to enrich mitotic cells. Cells with MAD2 positive chromosomes were counted (Mean and SEM, two-tailed Student’s t-test, n = 5 experiments; \*\**p* < 0.01). Scale bar, 20 μm. (H) Micronuclei observation in K562 WT or CENP-C^ΔM12BD^ cells. Arrowheads show typical micronuclei. Cells with micronuclei were counted (Mean and SEM, two-tailed Student’s t-test, n = 5 experiments; \*\**p* < 0.01). Scale bar, 20 μm.

**Figure S3.**
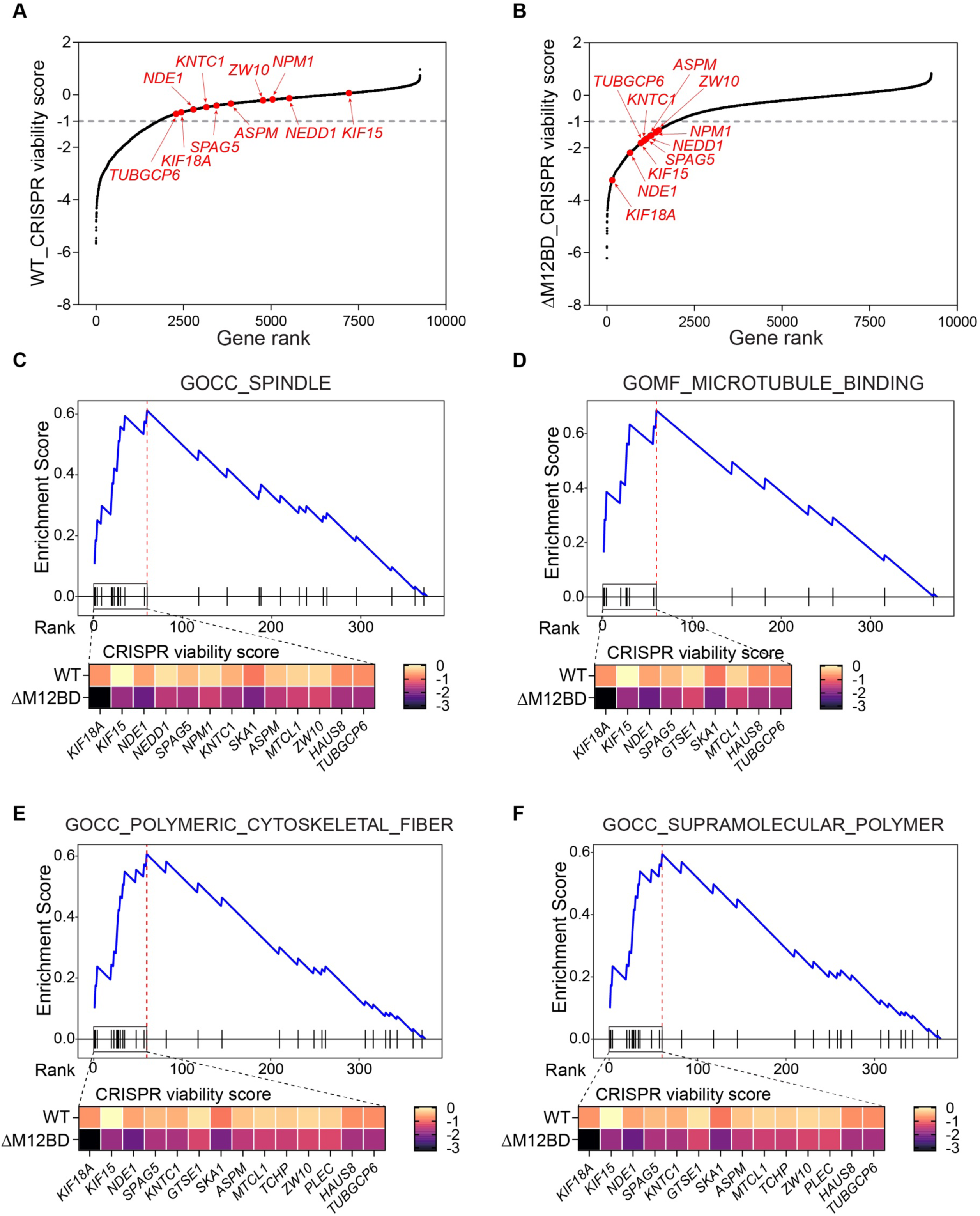
The CRISPR screening in K562 WT and CENP-C^ΔM12BD^ cells. (A) Distribution of CRISPR viability score of each gene in K562 WT cells. Gene with a median fold change (log2) > -1 was identified as a nonessential gene in K562 WT cells. (B) Distribution of CRISPR viability score of each gene in K562 CENP-C^ΔM12BD^ cells. Gene with a median fold change (log2) < -1 was identified as an essential gene in K562 CENP-C^ΔM12BD^ cells. (C-F) GSEA result of GOCC_SPINDLE gene set (C), GOMF_MICROTUBULE_BINDING gene set (D), GOCC_POLYMERIC_CYTOSKELETAL_FIBER gene set (E), and GOCC_SUPRAMOLECULAR_POLYMER gene set (F). The heat maps indicate the CRISPR viability score of leading edge genes of the preranking based on the magnitude of differences in CRISPR viability score between K562 WT and CENP-C**^Δ^**^M12BD^ cells, in descending order.

**Figure S4.**
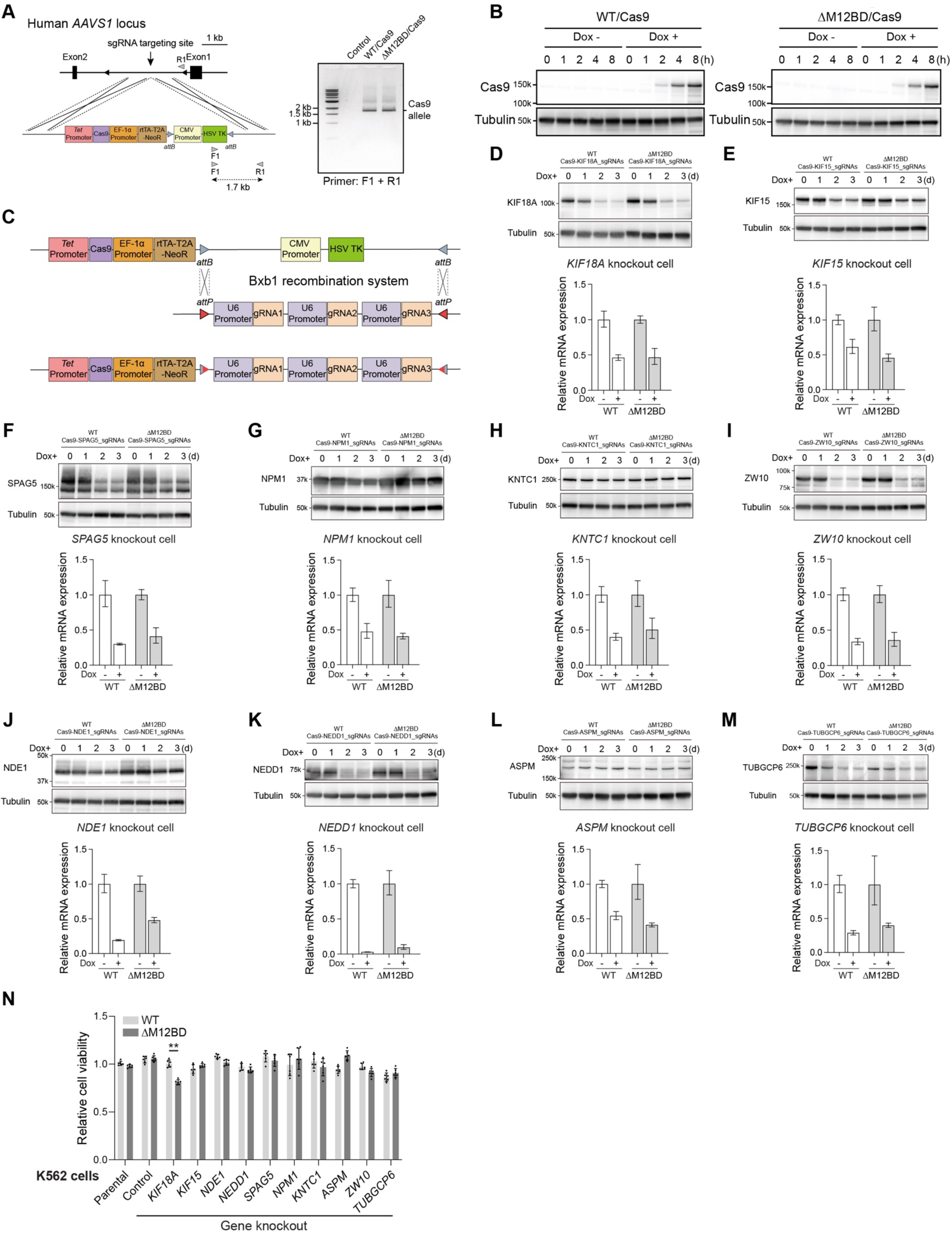
Generation of CRISPR-based conditional knockout cell lines for genes identified by the CRISPR screening in K562 WT and CENP-C^ΔM12BD^ cells. (A) Schematic representation of the strategy to insert the Cas9 construct under the control of Tet-responsible (Tet On) promoter into human *AAVS1* locus in K562 WT or CENP-C^ΔM12BD^ cells. The primers used for Genotyping PCR are shown. (B) Immunoblot analysis for conditional expression of Cas9 in K562 WT or CENP-C^ΔM12BD^ cells after Dox addition. (C) Strategy to insert sgRNAs for each target gene. As the Cas9 construct contains HSV TK flanked with two *attB* sequences, three U6 promoter-sgRNA sequences flanked by *attP* sequences can be inserted into this locus by *attB*-*attP* recombination with Bxb1 recombinase. (D-M) mRNA expression of each target gene in K562 WT and CENP-C^ΔM12BD^ cells after Dox addition. Expression of all tested genes was reduced at 48 h after Dox addition. mRNA expression was calculated by QuantStudio Design & Analysis Software v1.5.1 using the RQ (Relative Quantitation) value, error bars indicate RQ minimum and maximum (each sample size: n = 3). Immunoblot analyses of each protein in K562 WT and CENP-C^ΔM12BD^ cells at 0, 1, 2, and 3 days after Dox addition were also shown. KO efficiency for some proteins was not good. This may be due to the design of gRNAs. (N) Cell viability of K562 WT or CENP-C^ΔM12BD^ cells after knockout of indicated genes at 2 days after Dox addition (Mean and SD, two-tailed Student’s t-test, K562 WT cells: n = 6, K562 CENP-C^ΔM12BD^ cells: n = 6; \*\**p* < 0.01).

**Figure S5.**
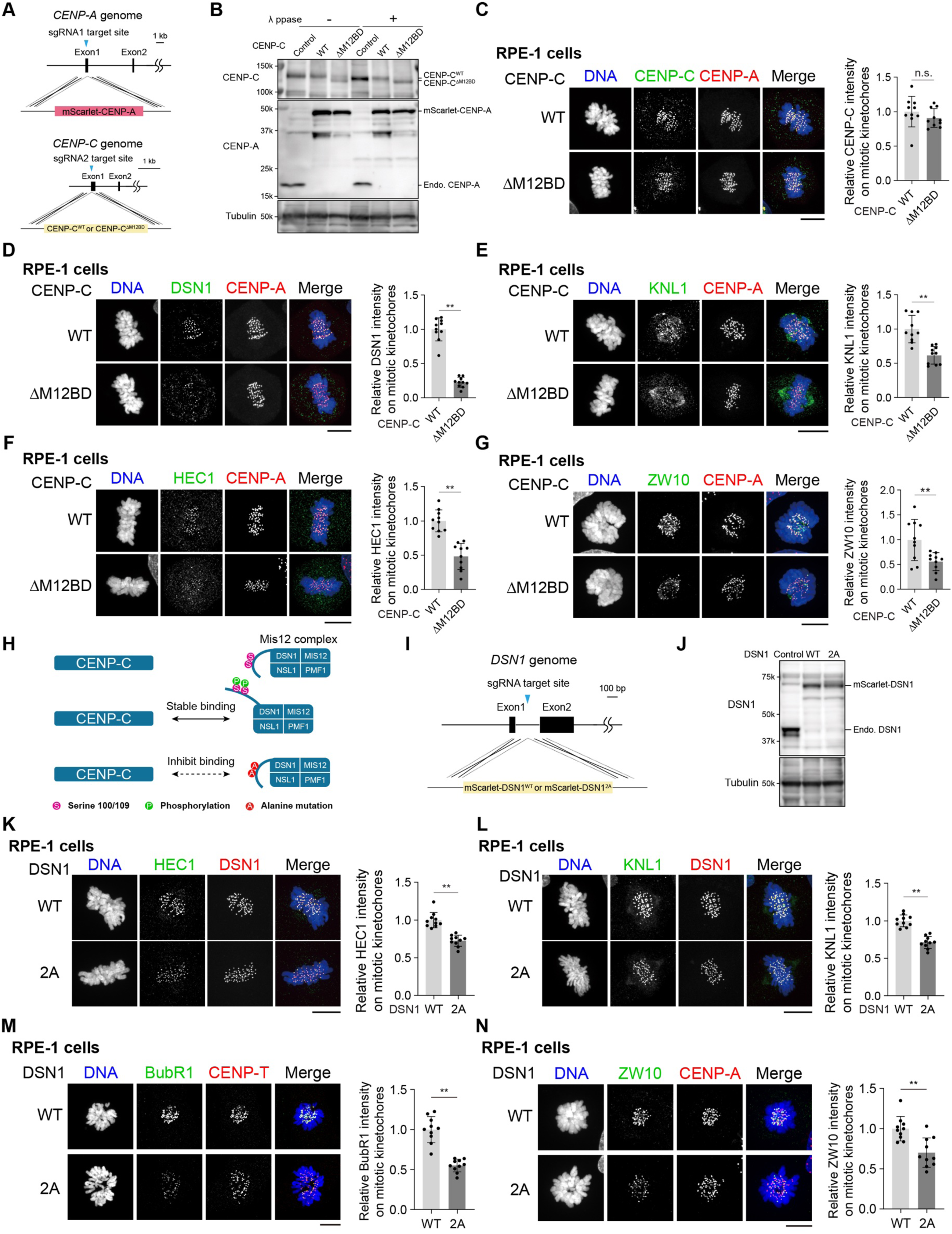
Generation and characterization of RPE-1 CENP-C ^WT^, CENP-C^ΔM12BD^ DSN1^WT^, and DSN1^2A^ cells. (A) Strategy to generate RPE-1 CENP-C^WT^ or CENP-C^ΔM12BD^ RPE-1 cells. mScarlet fused CENP-A was introduced into the endogenous *CENP-A* gene locus in RPE-1 cells. Then, either CENP-C^WT^ or CENP-C^ΔM12BD^ cDNA containing construct was integrated into the endogenous *CENP-C* gene locus in RPE-1 cells expressing mScarlet fused CENP-A. (B) Immunoblot analysis to detect CENP-C^WT^, CENP-C^ΔM12BD^, and mScarlet-CENP-A in each cell line. α-tubulin was probed as a loading control. λPPase was used to detect clear signals of CENP-C proteins as shown in K562 cells (λPPase +, Figure S1D). (C-G) CENP-C, DSN1, KNL1, HEC1, and ZW10 localization in RPE-1 CENP-C^WT^ or CENP-C^ΔM12BD^ cells. Each protein was stained with an antibody against each protein (green). mScarlet-CENP-A was used as a kinetochore marker (CENP-A, red). DNA was stained with DAPI (blue). Scale bar, 10 μm. Signal intensities of each protein at mitotic kinetochores were quantified (Mean and SD, two-tailed Student’s t-test, CENP-C^WT^ cells: n = 10 cells, CENP-C^ΔM12BD^ cells: n = 10 cells; n.s., non-significant; \*\**p* < 0.01). (H) Schematic representation of the regulation of CENP-C-Mis12C interaction. The basic motif of DSN1, a subunit of Mis12C, masks the CENP-C-binding surface of Mis12C, preventing the CENP-C-Mis12C interaction. Once Aurora B phosphorylates two residues (S100 and S109) on the basic motif, the mask is released, and the CENP-C-Mis12C interaction is facilitated. The alanine substitution of these sites reduces the CENP-C-Mis12C interaction. (I) Strategy to introduce mScarlet fused DSN1^WT^ or DSN1^2A^ cDNA into endogenous *DSN1* gene locus. The position of sgRNA target site is shown. (J) Immunoblot analysis to detect DSN1^WT^ or DSN1^2A^. α-tubulin was probed as a loading control. (K-N) HEC1 (K), KNL1 (L), BubR1 (M) and ZW10 (N) localization in RPE-1 DSN1^WT^ or DSN1^2A^ cells (green). mScarlet-DSN1 (either DSN1^WT^ or DSN1^2A^) was used as a kinetochore marker (red) for HEC1 and KNL1 staining. CENP-T or CENP-A was used as a kinetochore marker (red) for BubR1 and ZW10 staining, respectively. DNA was stained with DAPI (blue). Scale bar, 10 μm. Signal intensity of each protein at mitotic kinetochores was quantified (Mean and SD, two-tailed Student’s t-test, DSN1^WT^ cells: n = 10 cells, DSN1^2A^ cells: n = 10 cells; \*\**p* < 0.01).

**Figure S6.**
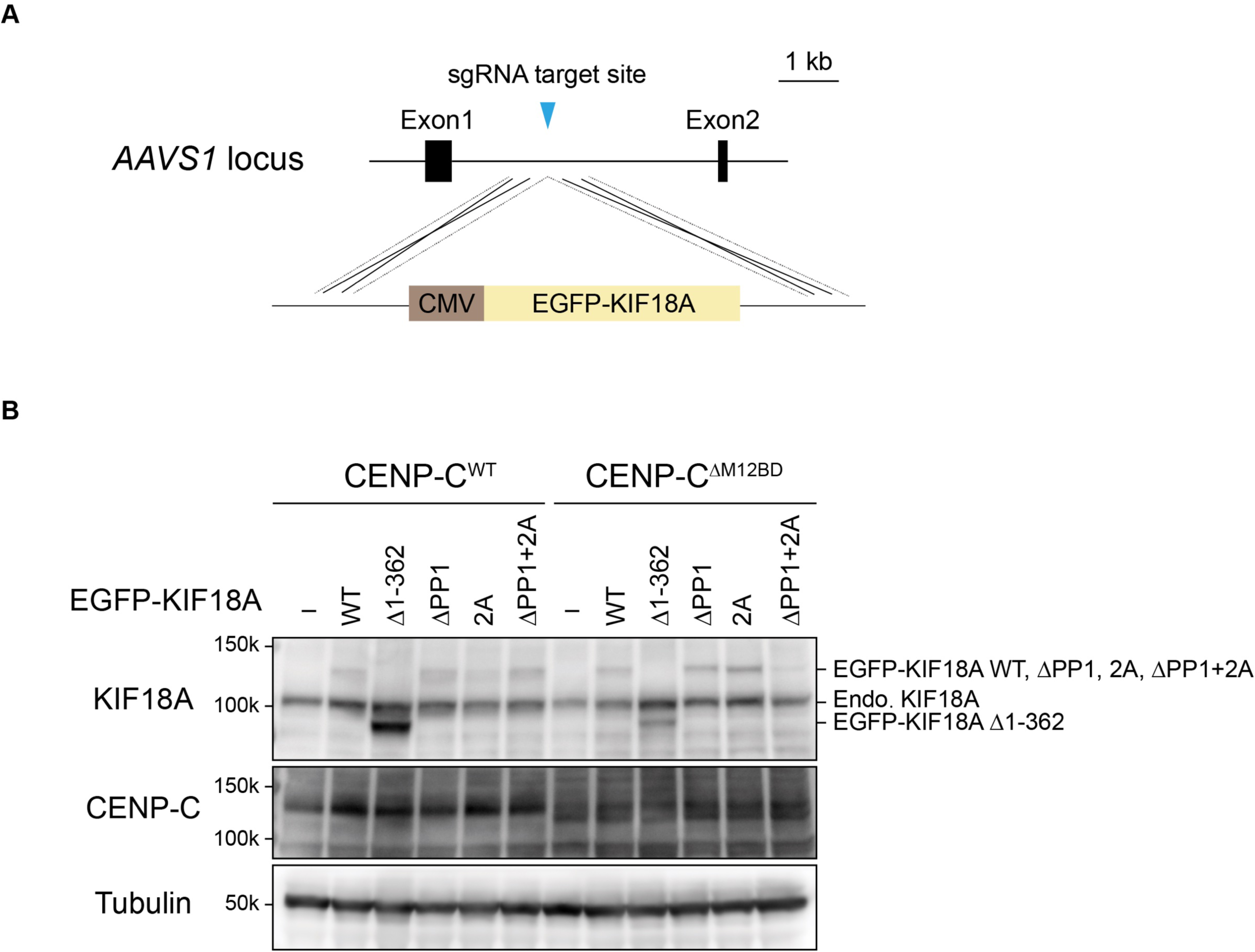
Integration of KIF18A mutants into *AAVS1* locus in RPE-1 cells (A) Strategy to insert EGFP-fused various versions of KIF18A cDNA into the *AAVS1* gene locus in RPE-1 CENP-C^WT^ or CENP-C^ΔM12BD^ cells. The position of the sgRNA target site is shown. (B) Immunoblot analysis to detect wild-type (WT) and various mutants of KIF18A in RPE-1 CENP-C^WT^ or CENP-C^ΔM12BD^ cells. α-tubulin was probed as a loading control.

**Figure S7.**
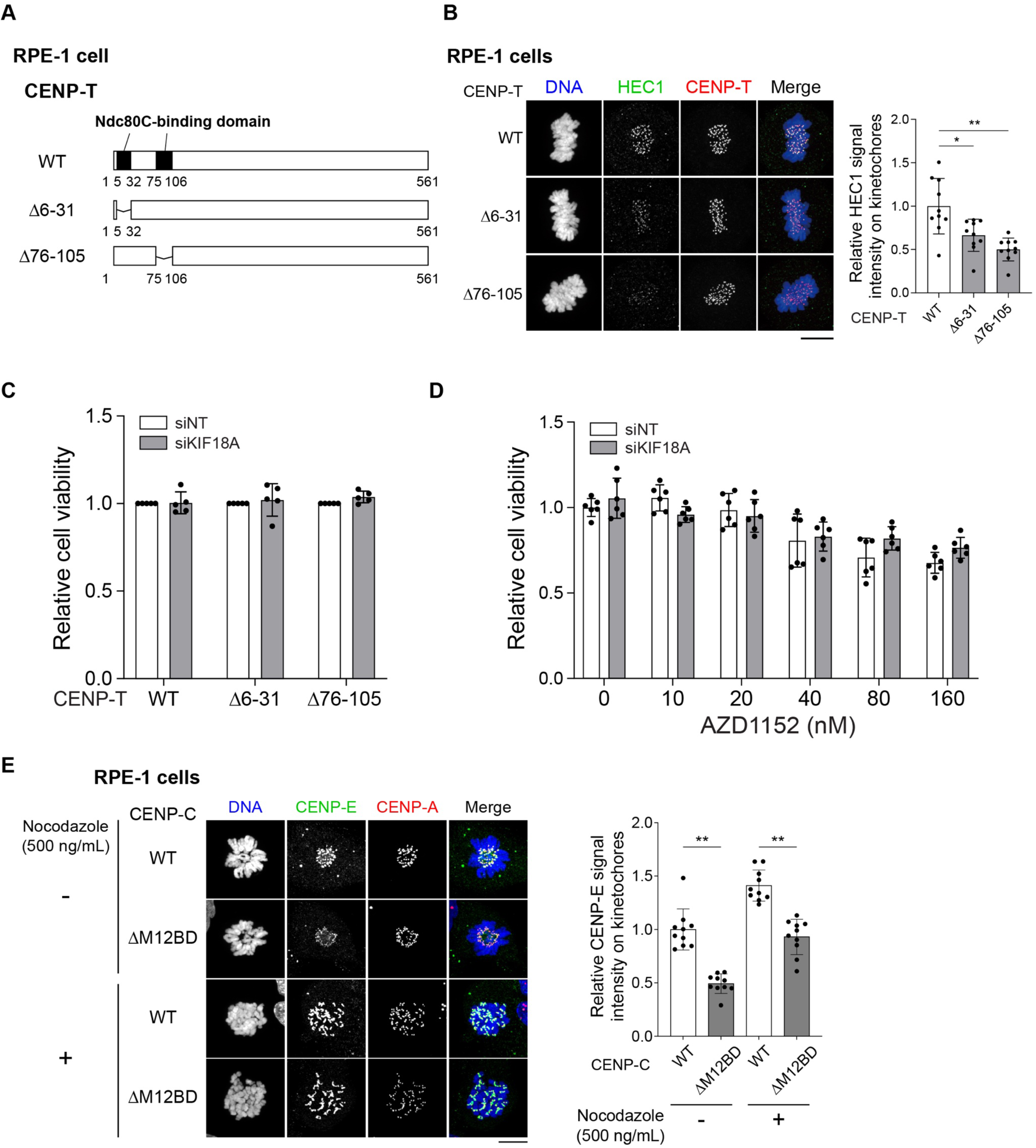
Neither CENP-T mutation lacking Ndc80C-binding domain nor inhibition of Aurora B activity shows synthetic lethality with *KIF18A* knockdown in RPE-1 cells. (A) Schematic representation of human CENP-T. Human wild-type CENP-T (WT) has two Ndc80C binding regions (amino acids 6-31 and 76-105). Each Ndc80C binding region was deleted in RPE-1 CENP-T^Δ6-31^ or CENP-T^Δ76-105^ cells, respectively. (B) HEC1 localization in RPE-1 CENP-T^Δ6-31^ or CENP-T^Δ76-105^ cells (green). CENP-T was used as a kinetochore marker (red). DNA was stained with DAPI (blue). Scale bar, 10 μm. Signal intensities of HEC1 at mitotic kinetochores in each cell line were quantified (Mean and SD, two-tailed Student’s t-test, CENP-T^WT^ cells: n = 10 cells, CENP-C^Δ6-31^ cells: n = 10 cells, CENP-C^Δ76-105^ cells: n = 10 cells; \**p* < 0.1; \*\**p* < 0.01). (C) Cell viability of RPE-1 CENP-T^WT^, CENP-T^Δ6-31^, or CENP-T^Δ6-31^ cells after treatment with siNT or siKIF18A. (Mean and SEM, two-tailed Student’s t-test, n = 5 independent experiments). (D) Cell viability of RPE-1 cells in the presence of various concentrations of AZD1152, an Aurora B inhibitor, with the treatment of siNT or siKIF18A. (Mean and SD, two-tailed Student’s t-test, each sample size: n = 6). (E) CENP-E localization in RPE-1 CENP-C^WT^ or CENP-C^ΔM12BD^ cells in the presence or absence of 500 ng/mL nocodazole treatment for 6 h. CENP-E was stained with an anti-CENP-E antibody (green). mScarlet-CENP-A was used as a kinetochore marker (CENP-A, red). DNA was stained with DAPI (blue). Scale bar, 10 μm. Signal intensity of CENP-E at mitotic kinetochores was quantified (Mean and SD, two-tailed Student’s t-test, CENP-C^WT^ cells: n = 10 cells, CENP-C^ΔM12BD^ cells: n = 10 cells; \*\**p* < 0.01).

**Table S1. CRISPR viability scores of tested genes in K562 WT and CENP-*C*^ΔM12BD^ cells**

**Table S2. sgRNA sequences used in this study**

**Table S3. PCR primer and siRNA sequences used in this study**

**Movie S1.** A time lapse movie of RPE1 CENP-C^WT^ cells after siNT treatment (Magenta: GFP-H2A; Yellow: mScarlet-CENP-A; Green: Tubulin).

**Movie S2.** A time lapse movie of RPE1 CENP-C^WT^ cells after siKIF18A treatment (Magenta: GFP-H2A; Yellow: mScarlet-CENP-A; Green: Tubulin).

**Movie S3.** A time lapse movie of RPE1 CENP-C^τιM12BD^ cells after siNT treatment (Magenta: GFP-H2A; Yellow: mScarlet-CENP-A; Green: Tubulin).

**Movie S4.** A time lapse movie of RPE1 CENP-C^τιM12BD^ cells after siKIF18A treatment (Magenta: GFP-H2A; Yellow: mScarlet-CENP-A; Green: Tubulin).

